# Widespread readthrough events in plants reveal unprecedented plasticity of stop codons

**DOI:** 10.1101/2023.03.20.533458

**Authors:** Yuqian Zhang, Hehuan Li, Yanting Shen, Shunxi Wang, Lei Tian, Haoqiang Yin, Jiawei Shi, Anqi Xing, Jinghua Zhang, Usman Ali, Abdul Sami, Xueyan Chen, Chenxuan Gao, Yangtao Zhao, Yajing Lyu, Xiaoxu Wang, Yanhui Chen, Zhixi Tian, Shu-Biao Wu, Liuji Wu

## Abstract

Stop codon readthrough (SCR), the decoding of a stop codon as a sense codon by the ribosome, has important biological implications but remains largely uncharacterized in plants. Here, we identified 1,009 SCR events in two monocots (maize, rice) and two dicots (soybean, *Arabidopsis*) using a proteogenomic strategy with 80 customized databases. SCR transcripts were mostly significantly shorter and had fewer components than non-SCR transcripts in two monocot plants, although these differences were not as significant in the dicots. Mass spectrometry evidence revealed that all three stop codons involved in SCR events could be recoded as 20 standard amino acids, some of which were also supported by suppressor transfer RNA analysis. In addition, we observed multiple functional signals in the C-terminal extensions of 34 maize SCR proteins, and characterized the structural and subcellular localization changes in the extended protein of BASIC TRANSCRIPTION FACTOR 3. Overall, our study not only demonstrates that SCR events are widespread in plants but also reveals the unprecedented recoding plasticity of stop codons, which provides important new insights into the flexibility of genetic decoding.

## Introduction

The rules of the genetic code were long considered universal dogma in all organisms after the standard genetic code was deciphered decades ago. However, exceptions to the rules have been increasingly observed in different species over the years, substantially altering the perception of the decoding rules from being static to dynamic^1, 2^. Living organisms are now widely understood to employ various modes to expand the coding potential of their genome, such as alternative translation initiation^3^, ribosome frameshifting^4^, and translational bypassing^5^. Stop codon readthrough (SCR) is another important means of recoding that is classified as a non-canonical translational event^6, 7^. SCR occurs when the ribosome reads a stop codon (UGA, UAG, or UAA) as a sense codon rather than the stop signal with the translation terminating at a later in-frame stop codon^7, 8^. Such an event can generate a C-terminally extended protein isoform which may acquire novel biological functions, since the amino acid sequence appended at the C-terminus possibly alters the activity^9^, stability^10^, and/or subcellular localization^2, 11^ of the protein. Indeed, SCR proteins have been implicated in a variety of biological processes, such as the regulation of intercellular cytoplasm flow^12^, neuronal integrity and reproductive health^13^, cellular redox homeostasis^2, 14^, and angiogenesis^15, 16^.

Initially discovered in viruses as a strategy to expand protein-coding potential^17–19^, SCR was subsequently identified in other organisms. In plant pathogens, *Phosphoglycerate kinase* (*PGK*) in *Ustilago maydis* and *Glyceraldehyde-3-phosphate dehydrogenase* (*GAPDH*) in *Aspergillus nidulans* are typical functional readthrough genes conserved in fungal species^2^. In the yeast *Saccharomyces cerevisiae*, the [PSI+] strain forms prion-like polymers of the abnormal eukaryotic release factor 3 (eRF3), which is implicated in translation termination and peptide release, resulting in a high level of readthrough across many genes^20^, including *Phosphodiesterase 2* (*PDE2*), *Interacting with mpp10p 3* (*IMP3*), and *Bypass of stop codon 4* (*BSC4*)^10, 21, 22^. In *Drosophila melanogaster*, readthrough is estimated to affect over 300 genes^23, 24^, such as *kelch* (*kel*)^12, 25^, *Synapsin* (*Syn*)^26^, *headcase* (*hdc*)^27^, and *traffic jam* (*tj*)^13^. In mammals, *Hemoglobin beta chain* (*HBB*)^28^, *Myelin protein zero* (*MPZ*)^29^, *Vascular endothelial growth factor-A* (*VEGF-A*)^15^, *Mitochondrial carrier 2* (*MTCH2*)^30^, *Argonaute 1* (*AGO1*)^31^, *Opioid related nociceptin receptor 1* (*OPRL1*)^32^, *Mitogen-activated protein kinase 10* (*MAPK10*)^32^, *Aquaporin 4* (*AQP4*)^33^, *Lactate dehydrogenase B* (*LDHB*)^34^, *Malate dehydrogenase 1* (*MDH1*)^34^, *Vitamin D receptor* (*VDR*)^35^, and *Adenosylmethionine decarboxylase 1* (*AMD1*)^36, 37^ have been confirmed to undergo readthrough. However, SCR events in plants remain largely uncharacterized, with only one study reporting 144 readthrough genes in *Arabidopsis* detected with ribosome profiling^38^. Therefore, further exploration of SCR events with a more robust analysis is warranted in a wide range of plant species.

It is crucial to identify the amino acids decoded at the stop codon sites, as they can strongly affect the functionality of readthrough protein isoforms^11, 21, 39^. Previous studies have shown that UGA could be translated to arginine, asparagine, cysteine, glutamine, glycine, serine, threonine, tryptophan, and tyrosine^30, 31, 39–41^; UAA to alanine, glutamate, glutamine, lysine, serine, tryptophan, and tyrosine^39, 42–45^; and UAG to alanine, arginine, aspartate, glutamate, glutamine, glycine, leucine, lysine, serine, threonine, tryptophan, tyrosine, and valine^30, 39, 46, 47^. However, knowledge about the amino acids translated at stop codon sites in SCR events remains unknown in Plantae. In addition, the potential for stop codons to be recoded as amino acids other than those reported above has not been investigated yet.

Here, we employed a mass spectrometry (MS)-based proteogenomic strategy to study SCR events in plants and to identify the amino acids at the readthrough stop codon sites. We detected a total of 1,009 SCR events in the monocots maize and rice, and dicots soybean and *Arabidopsis*, and characterized their features. Interestingly, we discovered that the three stop codons could be recoded as all 20 standard amino acids, which reveals the unprecedented recoding plasticity of stop codons and the dynamic nature of genetic decoding. We also delineated the first comprehensive landscape of SCR events in plants at the protein level, uncovering a collection of previously unknown readthrough proteins to further explore their biological functions.

## Results

### A proteogenomic strategy to identify SCR events in plants

The SCR event is a means of alternative translation by redefining UGA, UAG, or UAA to specify amino acids, thus extending the translation at the C-terminus. The most definitive evidence for SCR occurrence is the direct detection of translation at the stop codon sites. However, SCR events are non-canonical and cannot be identified by a typical proteomic strategy. Therefore, we implemented a proteogenomic strategy integrating MS-based proteomics with 20 customized databases to identify SCR events in maize at the protein level (Fig. 1). First, total protein was extracted from maize leaves, followed by digestion, fractionation, and liquid chromatography-tandem mass spectrometry (LC-MS/MS) analysis to obtain MS/MS datasets of peptides for subsequent identification. We then constructed 20 customized databases with in-house scripts as described in the Methods section.

**Fig. 1.**
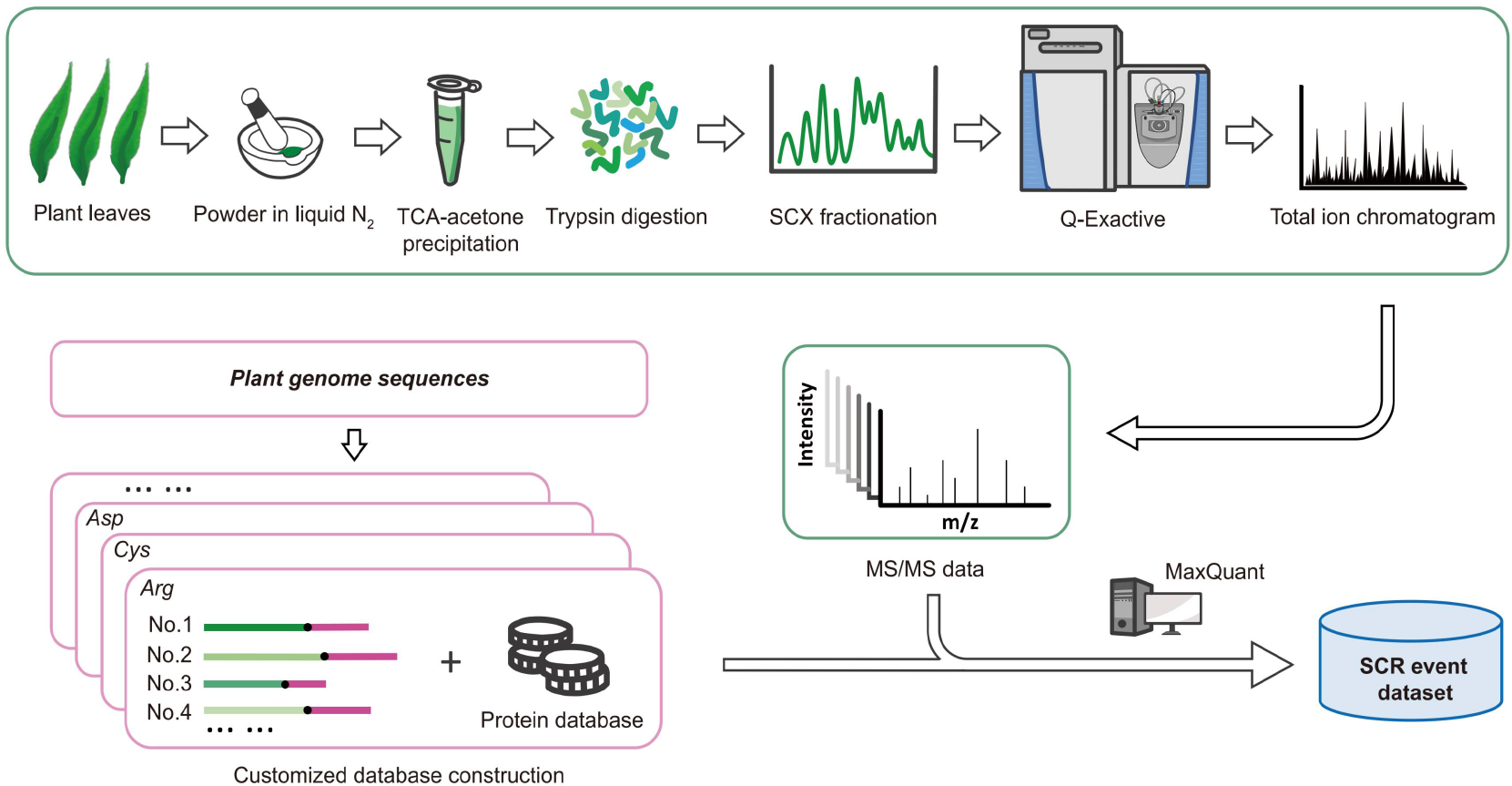
Proteogenomic workflow for identifying plant stop codon readthrough (SCR) events. The strategy consists of two procedures: shotgun proteomics experiment (green box) and customized database construction (pink boxes). Total protein was extracted from plant leaves and subjected to digestion, fragmentation, and LC-MS/MS analysis for identification of spectra datasets. Customized databases were constructed as described in Methods. MS/MS spectra datasets were searched against the 20 customized databases merged with the public plant protein database by the MaxQuant. We defined the MS/MS-identified peptides that exclusively spanned the annotated stop codon sites as “through” peptides, and classified proteins sharing a unique stop codon site and containing a “through” peptide as one SCR event.

To capture SCR events in maize, we used the obtained MS/MS data to search against the 20 SCR customized databases with MaxQuant search engine. This strategy identified 708 unique “through” peptides spanning the stop codon site. For further quality control of MS data, we next applied more stringent criteria for “through” peptides to identify SCR events with high confidence (see Methods). The screening steps resulted in much more reliable “through” peptides, which corresponded to 367 SCR events involving 1,063 proteins (Supplementary Table 1). All identified peptides for SCR proteins are listed in Supplementary Table 2. Collectively, these results not only demonstrate the abundance of readthrough events naturally occurring in maize, but also uncover a previously unknown source of protein isoforms produced by SCR.

### Verifying SCR events in maize

To verify the SCR events in maize identified above, we retrieved the available ribosome profiling (Ribo-seq) datasets of 56 maize samples from NCBI to analyze the SCR events at the mRNA level. Four Ribo-seq datasets with higher quality were finally retained after further screening (see Methods). We then analyzed the three-nucleotide (3-nt) periodicity of the Ribo-seq reads (30-mers) in the coding sequences (CDSs), putative extensions, and distal 3′ untranslated regions (UTRs). The results showed that the putative extensions exhibited a visible hallmark of 3-nt periodicity (*P <* 0.05, Fig. 2a), i.e., the reading frame 0 occurred significantly higher than the other two, which was similar to that of the CDSs (Fig. 2a). By contrast, such difference was not observed in the distal 3′ UTRs (Fig. 2a). In addition, the average ribosome footprint density (in RPKM) of putative extensions was significantly higher than that of the distal 3′ UTRs in the four datasets (*P <* 0.001, Fig. 2b). These results suggest that readthrough is a significant contributor to the ribosome footprint density in putative extensions. After applying the conditions described in the Methods section for the four Ribo-seq datasets to verify SCR, we found that 55 out of the 367 SCR events identified by the proteogenomic approach were also corroborated by Ribo-seq analysis (Fig. 2c and Supplementary Table 3).

**Fig. 2.**
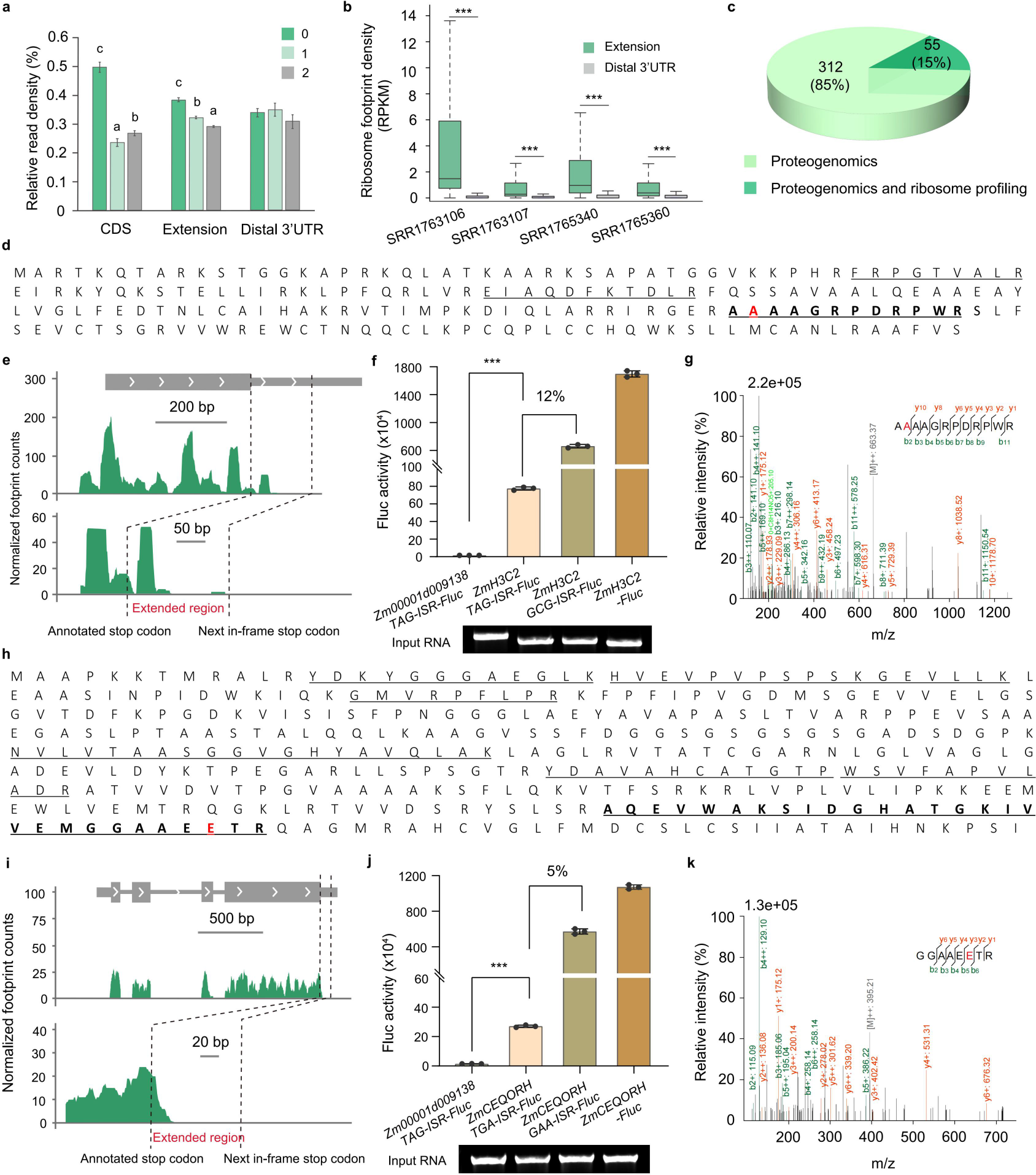
Verifying SCR events in maize. **a** Three-nucleotide periodicity of 30-mers in the CDSs, putative extensions, and the distal 3′ UTRs. Data are mean ± s.d. Different letters indicate significant differences at *P* < 0.05 by one-way ANOVA with Tukey’s multiple comparisons test. Green, light green, and gray bars represent frame 0, 1, and 2, respectively. **b** Distribution of ribosome footprint density in putative extensions and the distal 3′ UTRs in the four Ribo-seq datasets. Distal 3′ UTR was defined as the region 40 codons after the next stop-codon or the region from the next stop-codon to the third stop codon within 40 codons. Box edges represent the 0.25 and 0.75 quantiles. The bold lines represent the median and whiskers represent 1.5×IQR. Statistical significance was determined using a two-sided Student’s *t*-test, \*\*\**P* < 0.001. **c** The proportion of SCR events confirmed by ribosome profiling among those identified by proteogenomics. **d-k** Validation of two readthrough examples, *ZmH3C2* (**d-g**) and *ZmCEQORH* (**h-k**). **d, h** The complete protein sequences extended to the next in-frame stop codon of the two genes showing the underlined peptides identified by mass spectrometry analysis, and the readthrough peptides are in bold. The amino acids translated at the stop codon sites in readthrough peptides are highlighted in red, which are alanine (A) at UAG (**d**) and glutamate (E) at UGA (**h**). **e, i** Ribo-seq analysis revealing clear footprint signals in the C-terminal extensions of the two transcripts. Dashed lines represent the annotated stop codon and the next in-frame stop codon. **f, j** *In vitro* translation experiment showing readthrough activities of *ZmH3C2* and *ZmCEQORH*. Plasmids containing test cDNA-stop codon-ISR-*Fluc* were firstly transcribed *in vitro*, followed by *in vitro* translation using wheat germ extract. Luciferase activities were used to represent the readthrough activities. Constructs with the stop codon mutated to a sense codon were used as a positive control of the translation to measure the efficiency of SCR. The non-SCR gene *Zm00001d009138* with its TAG and ISR sequence fused with *Fluc* (*Zm00001d009138-TAG-ISR-Fluc*) was used as the SCR-negative control, as it showed no SCR sign in either proteogenomics or Ribo-seq analysis. All samples were tested in biological triplicates. Data are mean ± s.e.m (n=3). Asterisks indicate statistical significance by two-sided Student’s *t*-test (\*\*\**P* < 0.001). ISR, inter-stop codon region (extension region). **g, k** MS/MS spectra of the identified readthrough peptides AAAAGRPDRPWR for *ZmH3C2* (**g**) and GGAAEETR for *ZmCEQORH* **(k)** readthrough products expressed from (**f**) and (**j**), respectively. Source data are provided as a Source Data file.

We selected two SCR representative genes, *ZmH3C2* and *ZmCEQORH*, which exerted functional roles in distinct biological processes related to chromatin assembly and stress response, respectively, for further analysis. For *ZmH3C2*, which encodes HISTONE H3.2, MS analysis identified the readthrough peptide AAAAGRPDRPWR with its second alanine (A) translated at the UAG site (Fig. 2d and Supplementary Fig. 1a). The readthrough of this gene was also evidenced by ribosome footprints within the extended region downstream of the annotated stop codon (Fig. 2e). For *ZmCEQORH*, encoding CHLOROPLAST ENVELOPE QUINONE OXIDOREDUCTASE, MS analysis identified the “through” peptide AQEVWAKSIDGHATGKIVVEMGGAAEETR, in which the antepenultimate glutamate (E) was incorporated at the UGA site (Fig. 2h and Supplementary Fig. 1b). We also observed ribosome footprints beyond the annotated stop codon of the corresponding transcript (Fig. 2i).

In addition, we performed *in vitro* translation experiments using wheat germ extract to further demonstrate the readthrough of the above two genes. The results showed that *ZmH3C2* and *ZmCEQORH* had significantly higher readthrough activities compared to the SCR-negative gene *Zm00001d009138* (*P* < 0.001), as demonstrated by the detection of luciferase activities. The SCR rates of *ZmH3C2* and *ZmCEQORH* were 12% and 5%, respectively (Fig. 2f, j). It is known that the rate of the basal translation error is lower than 0.1%^8^, thus the results indicate that *ZmH3C2* and *ZmCEQORH* underwent the programmed SCR events. Further MS analysis of the readthrough products identified the “through” peptides AAAAGRPDRPWR and GGAAEETR for the two genes, where the UAG and UGA were recoded as alanine and glutamate (Fig. 2g, k), respectively. Taken together, our Ribo-seq analysis, *in vitro* translation experiments, and MS analysis validated the SCR events identified using proteogenomics in maize, and demonstrated the feasibility of the proteogenomic method for large-scale identification of SCR events in plants.

### Characteristics of SCR events in maize

To understand the characteristics of SCR events in maize, we first analyzed the stop codon preferences. All three stop codons were read through among the 367 SCR events: UGA was involved in 174 events (47.4%), followed by UAG in 102 events (27.8%) and UAA in the remaining 91 events (24.8%) (Fig. 3a). We found that UGA was the most frequent next in-frame stop codon following these readthrough stop codons, accounting for 76 cases after the readthrough UGA, 46 cases after the readthrough UAG, and 40 cases after the readthrough UAA (Fig. 3a). The genome-wide distribution pattern showed that these SCR events were distributed heterogeneously across each chromosome of maize, and many of the SCR events were closer to the telomeres (Fig. 3b). In addition, a collection of 379 hotspot regions were obtained, 244 of which were SCR hotspot regions while 135 were non-SCR hotspot regions (Fig. 3b). These results revealed the wide distribution of SCR events in the maize genome.

**Fig. 3.**
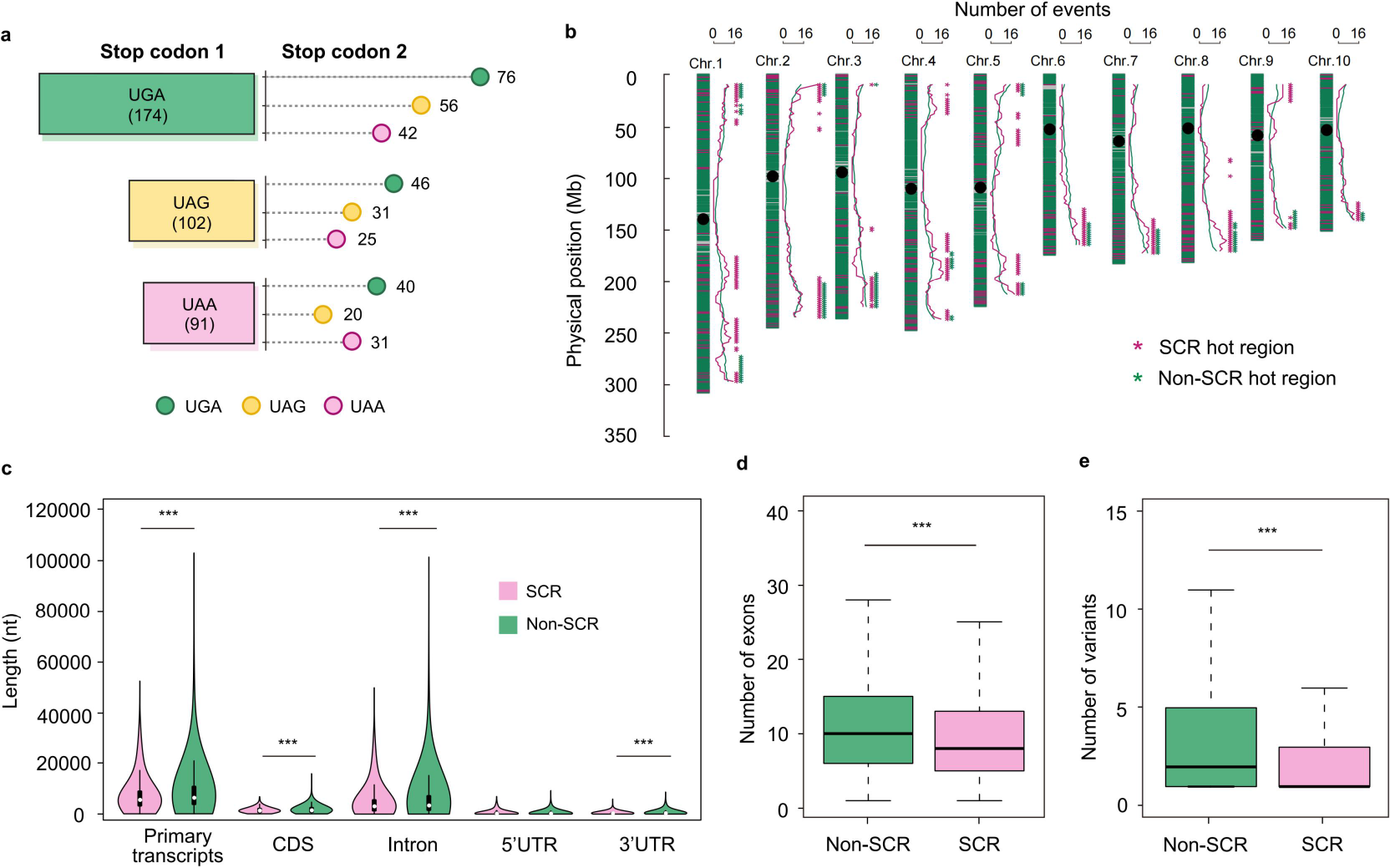
Characteristics of SCR events in maize. **a** Stop codon usage and the next stop codon preference in SCR events. Stop codon 1, readthrough stop codon; Stop codon 2, the next in-frame stop codon. **b** Genome-wide distribution of SCR (pink) and non-SCR (green) events in maize. For each chromosome, the distribution contains three columns. Left: SCR (pink) and non-SCR (green) events across the chromosomes. Black dots are centromeres. Middle: distribution patterns of SCR (pink) and non-SCR (green) events are shown by curves, shown as a 20-Mb sliding window with a 2-Mb step based on the B73 reference genome. Right: distribution of hotspot regions for SCR (pink) and non-SCR (green) events shown with asterisks. We defined hotspot regions as ≥5 SCR events and ≥335 non-SCR events in a 20-Mb window. **c** Violin plot describing the length distribution of primary transcripts and transcript elements in SCR and non-SCR events. The length of the primary transcript is the sum of CDSs, introns, and UTRs. CDS length represents the length of the open reading frame (ORF). Internal boxplots show the median values (white dots in the center) and the interquartile range (IQR; 0.25 and 0.75 quantiles) of the length distribution. **d** Number of exons in SCR and non-SCR primary transcripts. **e** Number of splice variants in SCR and non-SCR primary transcripts. For all boxplots, box edges represent the 0.25 and 0.75 quantiles. The bold lines represent the median values and the whiskers represent 1.5×IQR. \*\*\**P* < 0.001 (two-sided Student’s *t*-test). Source data are provided as a Source Data file.

Furthermore, we compared the length and complexity of SCR and non-SCR transcripts. Interestingly, we observed that the SCR primary transcripts were significantly shorter than the non-SCR transcripts (Fig. 3c). The average lengths of the elements from the primary transcripts, including coding sequences (CDSs), introns, and 3′ UTRs, were also markedly shorter in SCR events than those in non-SCR events (Fig. 3c). The 5′ UTR was also shorter in readthrough transcripts, albeit not statistically significant. Consistent with the length of readthrough transcripts, their corresponding proteins had notably shorter lengths with fewer amino acids (Supplementary Fig. 2a) and lower molecular weight (Supplementary Fig. 2b). Moreover, SCR primary transcripts had significantly fewer exons (Fig. 3d) and splice variants (Fig. 3e) than those of non-SCR primary transcripts.

Earlier studies have shown that *cis*-acting elements such as nucleotides immediately surrounding the readthrough stop codon and motifs downstream the stop codon can influence readthrough^48–50^. We thus examined the local mRNA context of SCR transcripts. An analysis of nucleotide frequencies in positions adjacent to the readthrough stop codons showed a bias of A>U>C>G at the -2 position, A>G>C>U at the -1 position, A>G>U>C at the +4 position, and U>C>G>A at the +5 position (Supplementary Fig. 3a). The most frequent tetranucleotide pattern (stop codon plus the following nucleotide) was UGA-U (Supplementary Fig. 3b). Extended regions downstream of the annotated stop codons in the readthrough transcripts were enriched with featured sequence motifs (Supplementary Dataset 1). Three motif examples, ZNF263, ASCL1, and ETV6 are shown in Supplementary Fig. 3c. These motifs enriched in extended regions likely exert effects on the particular facets of SCR by interacting with specific proteins. In addition, extended regions of SCR transcripts harbored remarkably lower free energy than their analogs of non-SCR transcripts (Supplementary Fig. 3d), suggesting more stable secondary structures in the extensions of the readthrough transcripts.

### Stop codons are recoded as 20 amino acids in maize SCR events

Since translation of the stop codon is crucial for SCR events, we examined which amino acids were translated at every readthrough stop codon site based on the MS data. Quite unexpectedly, we found that, with the exception of UAA which was recoded to 19 amino acids (except serine), UGA and UAG were translated to all 20 standard amino acids in maize SCR events (Fig. 4a). Among all the amino acids, glutamate was the most recoded amino acid for UGA (in 17 SCR events, 8.9%), methionine for UAG (in 17 cases, 15.6%), and aspartate for UAA (in 11 cases, 11.6%).

**Fig. 4.**
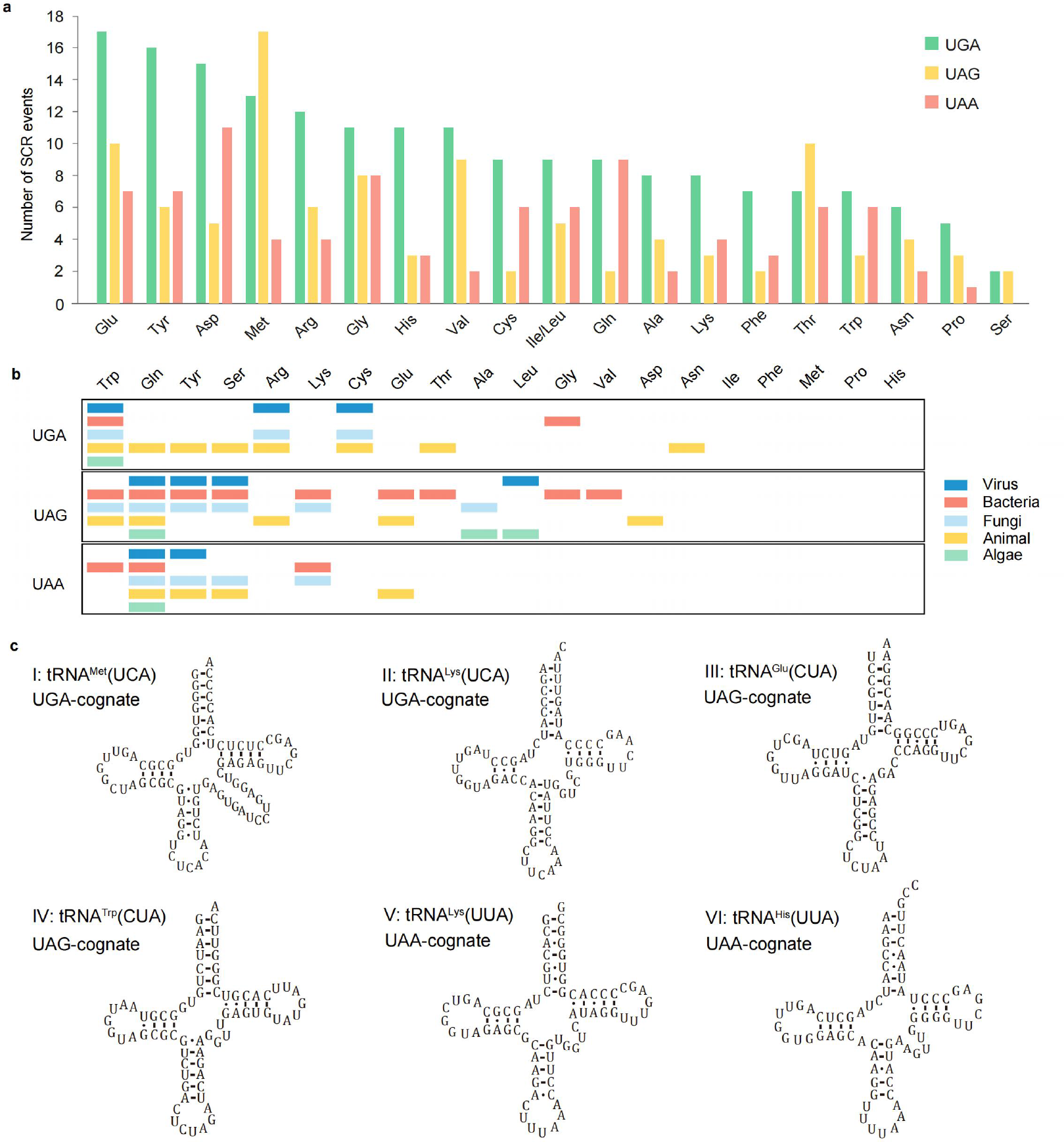
Stop codons are recoded as 20 amino acids in maize SCR events. **a** Frequency of amino acids specified by the three stop codons. The x-axis represents the 20 standard amino acids. The y-axis is the number of SCR events. **b** Previously reported amino acids translated at UGA, UAG, and UAA in different species. Blue, virus; orange, bacteria; light blue, fungi; yellow, animal; green, algae. **c** Examples of UGA-, UAG-, and UAA suppressor tRNAs identified by tRNAscan-SE and ARAGORN corresponding to amino acids shown at UGA, UAG, and UAA. The predicted secondary structures of the tRNAs are shown (I-VI): tRNA^Met^ (UCA) and tRNA^Lys^ (UCA) recognize UGA, tRNA^Glu^ (CUA) and tRNA^Trp^ (CUA) recognize UAG, and tRNA^Lys^ (UUA) and tRNA^His^ (UUA) recognize UAA.

Previous literature has reported several amino acids that were able to be translated at stop codon sites in different species (Fig. 4b and Supplementary Table 5), prompting us to explore the reasons behind the recoding plasticity of the stop codons in our results. Suppressor transfer RNA s (tRNAs) are specific tRNA species forming classical base pairing with the stop codon, which can lead to the translation^24^. We thus conducted a genome-wide search for such potential suppressor tRNAs capable of reading stop codons in maize by running tRNAscan-SE^51^ and ARAGORN^52^. As a result, we acquired a collection of UGA-, UAG-, and UAA-recognizing tRNAs. Multiple sequence alignments of the predicted suppressor tRNAs against the NCBI Nucleotide (NT) database resulted in different groups of suppressor tRNAs well aligned with the canonical tRNAs of a range of amino acids (Supplementary Table 6). We identified six classes of UGA suppressor tRNAs (Supplementary Tables 5 and 6): tRNA^Met^(UCA), tRNA^Lys^(UCA), tRNA^Val^(UCA), tRNA^Gly^(UCA), tRNA^Ile^(UCA), and tRNA^Thr^(UCA). The predicted secondary structures of the first two UGA suppressor tRNAs are shown in Fig. 4c I, II, and their nucleotide sequence alignments are presented in Supplementary Fig. 4a, b. We also detected six classes of UAG suppressor tRNAs (Supplementary Tables 5 and 6): tRNA^Glu^(CUA) (Fig. 4c III and Supplementary Fig. 5a), tRNA^Trp^(CUA) (Fig. 4c IV and Supplementary Fig. 5b), tRNA^Arg^(CUA), tRNA^Asp^(CUA), tRNA^Gly^(CUA), and tRNA^Lys^(CUA). Furthermore, we identified eight classes of UAA suppressor tRNAs (Supplementary Tables 5 and 6): tRNA^Lys^(UUA) (Fig. 4c V and Supplementary Fig. 6a), tRNA^His^(UUA) (Fig. 4c VI and Supplementary Fig. 6b), tRNA^Ala^(UUA), tRNA^Asn^(UUA), tRNA^Gly^(UUA), tRNA^Phe^(UUA), tRNA^Met^(UUA), and tRNA^Thr^(UUA). Collectively, the suppressor tRNAs identified in maize provide potential reasons for the translation of at least some amino acids at stop codon sites.

### SCR events and recoding plasticity of stop codons in other plants

In light of the abundant SCR events identified in maize, we questioned if SCR also occurred in other plant species. We thus followed the same proteogenomic method with the respective 20 custom databases to explore SCR events in rice, soybean, and *Arabidopsis*. Indeed, we detected abundant SCR events in these species with similar features to those in maize. We identified 198 SCR events in rice (Supplementary Table 7), 179 SCR events in soybean (Supplementary Table 9), and 265 SCR events in *Arabidopsis* (Supplementary Table 11), corresponding to 269, 385, and 468 proteins, respectively.

In rice, we observed that the stop codon usage SCR events was in the order of UGA > UAA > UAG, and the next in-frame stop codon following the readthrough stop codons also displayed a preference for UGA (Fig. 5a). For the dicots soybean and *Arabidopsis*, the stop codon usages were the same in the order of UGA > UAA > UAG in SCR events, while the most frequent next in-frame stop codon after the readthrough stop codons was UGA for soybean and UAA for *Arabidopsis* (Fig. 5b, c). Genome-wide distribution analysis showed that SCR events were distributed across all chromosomes in their respective plant species (Fig. 5d, e, f). We also analyzed the features of SCR and non-SCR transcripts in the three plants. In rice, the length of SCR primary transcripts and transcript elements including CDSs, introns and the 5′ UTR were shorter than those of non-SCRs (*P* < 0.05, Supplementary Fig. 7a, b). In addition, the SCR primary transcripts tended to possess fewer exons (Supplementary Fig. 7c) and had significantly fewer splice variants (*P* < 0.001, Supplementary Fig. 7d) than non-SCR transcripts. In the dicots soybean and *Arabidopsis*, however, the SCR transcript features were not apparent as compared to those in monocots. SCR transcripts in soybean had significantly shorter UTR lengths and fewer exons than the non-SCR transcripts (*P* < 0.05, Supplementary Fig. 8b, c) but no significance for other feature comparison (Supplementary Fig. 8a, b, d). Similarly, SCR transcripts from *Arabidopsis* showed no significant different from the non-SCR counterparts except fewer splice variants (Supplementary Fig. 9a, b, c, d).

**Fig. 5.**
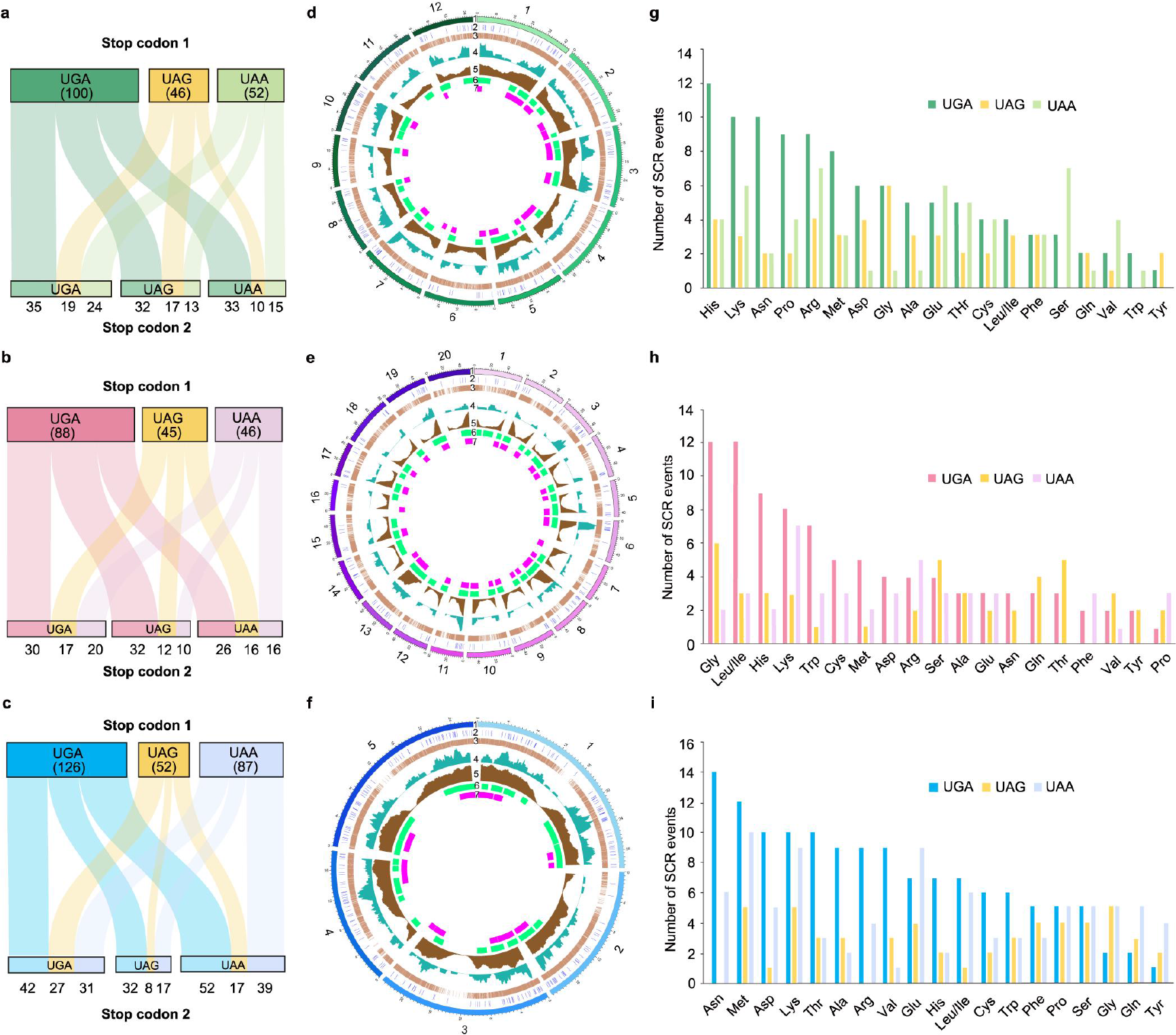
SCR events and recoding plasticity of stop codons in other plants. **a, b, c** Stop codon usage and the next stop codon preference of SCR events in rice (**a**), soybean (**b**), and *Arabidopsis* (**c**). Stop codon 1, readthrough stop codon; Stop codon 2, the next in-frame stop codon. **d, e, f** Circos plots showing the genome-wide distribution of SCR and non-SCR events in rice (**d**), soybean (**e**), and *Arabidopsis* (**f**). Each Circos plot consists of seven layers. 1, chromosomes; 2-3, distribution patterns of SCR (2) and non-SCR (3) events across the chromosomes; 4-5, distribution patterns of SCR (4) and non-SCR (5) events determined using a window size of 4 Mb with a step of 400 kb based on the Nipponbare reference genome for rice, a 10-Mb window with a 1-Mb step based on the ZH13 reference genome, and a 1.2-Mb window with a 100-kb step based on the Columbia-0 reference genome; 6-7, distribution of hotspot regions for SCR (6) and non-SCR (7) events. Hotspot regions were defined as ≥3 SCR events and ≥178 non-SCR events in a 4-Mb window for rice, ≥2 SCR events and ≥253 non-SCR events in a 10-Mb window for soybean, and ≥4 SCR events and ≥157 for non-SCR events in a 1.2-Mb window for *Arabidopsis*. **g, h, i** Frequency of amino acids specified by the three stop codons in rice (**g**), soybean (**h**), and *Arabidopsis* (**i**) SCR events. The x-axis represents the 20 standard amino acids. The y-axis is the number of SCR events.

Analysis of the recoded amino acids at stop codon sites revealed that UGA exhibited great plasticity to be recoded as the 20 standard amino acids in all the three plant species (Fig. 5g, h, i). For UAG, it was recoded to 18 amino acids (except serine and tryptophan) in rice (Fig. 5g), 17 amino acids (except cystine, aspartate, and phenylalanine) in soybean (Fig. 5h), and 18 amino acids (except aspartate and arginine) in *Arabidopsis* (Fig. 5i). Similarly, UAA was translated to 17, 16, and 20 amino acids in rice, soybean, and *Arabidopsis*, respectively (Fig. 5g, h, i). These results further uncover the high recoding plasticity of stop codons in plant SCR events.

### C-terminal extensions in maize SCR events contain multiple functional signals

To examine the biological implications of SCR events, we searched the functional signals in all C-terminal extensions of readthrough proteins in maize. As a result, multiple functional signals were detected in 34 extensions, including the transmembrane domains in 28 extensions, nuclear localization signals in two extensions, and peroxisomal targeting signals (PTS1) in four extensions (Fig. 6a and Supplementary Table 13). Among them, we selected BASIC TRANSCRIPTION FACTOR 3 (ZmBTF3), which was predicted to possess a transmembrane domain in its C-terminal extension (54 amino acids), for further analysis. First, we conducted the *in vitro* translation experiment to validate the readthrough of this gene. We observed high readthrough activity of *ZmBTF3* with an SCR rate of 15% (Fig. 6b), which revealed that *ZmBTF3* was read through and generated a novel isoform (hereafter termed ZmBTF3x). Next, we confirmed the expression of the fusion protein ZmBTF3x-Fluc by immunoblotting using an anti-Fluc antibody (Fig. 6c). The results showed that the molecular weight of the readthrough protein ZmBTF3x-Fluc was the same as its positive control ZmBTF3x^ACA^-Fluc and higher than that of ZmBTF3-Fluc (Fig. 6c). By contrast, no visible band of readthrough products expressed from the SCR-negative control *Zm00001d009138-TAG-ISR-Fluc* was observed in the immunoblot (Fig. 6c). The fusion protein ZmBTF3x-Fluc was then subjected to mass spectrometric analysis, resulting in the detection of different types of peptides labeled as “before”, “through”, and “after”, which further validated the production of the readthrough protein isoform ZmBTF3x (Fig. 6d, top). The spectrum for “through” peptide “EPEEKKESTFCCV” revealed that the stop codon UAA was translated as threonine (T) (Fig. 6d, bottom).

**Fig. 6.**
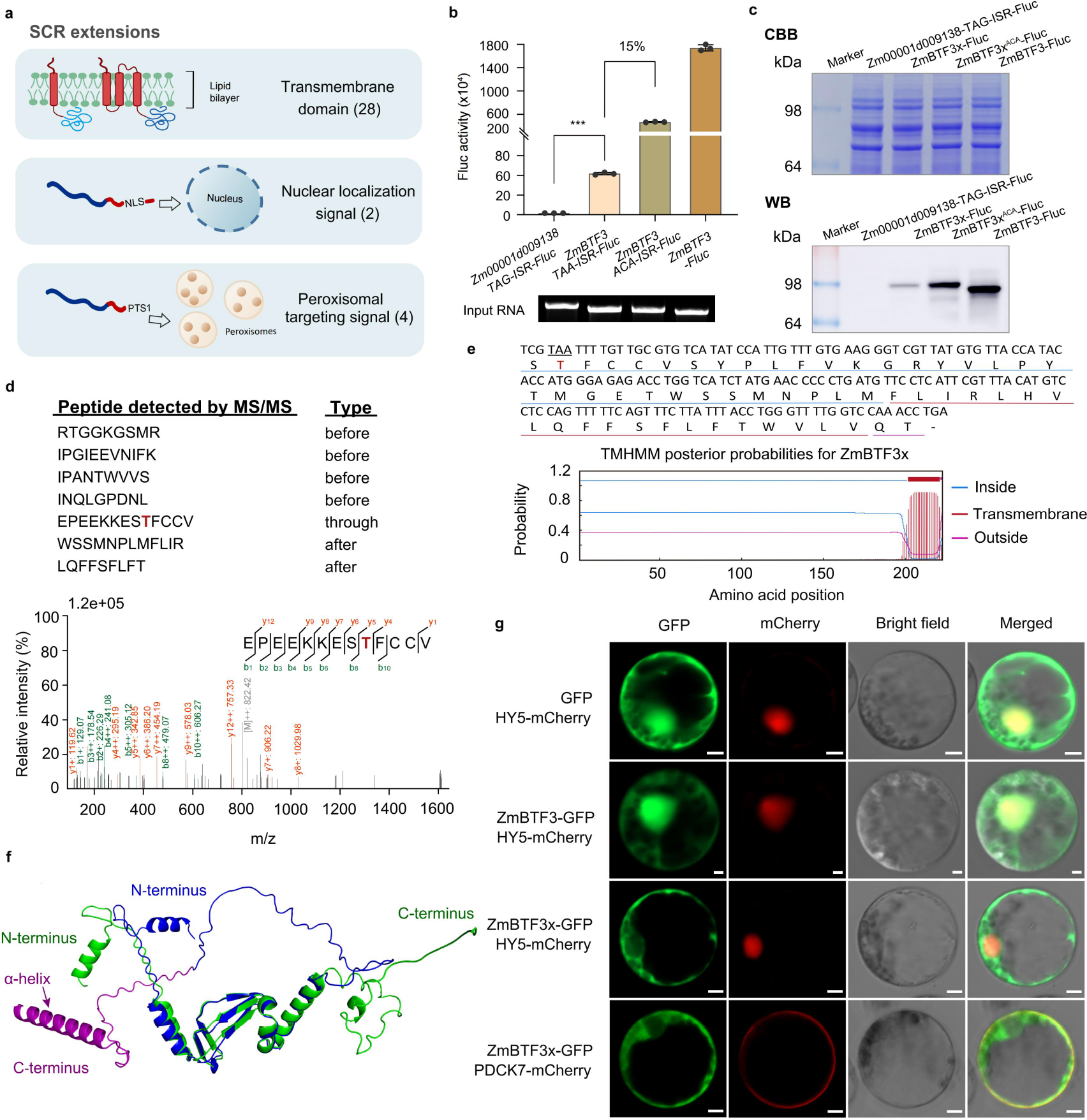
C-terminal extensions in maize SCR events contain multiple functional signals. **a** Multiple functional signals predicted in the extensions of SCR events. **b** Readthrough validation of *ZmBTF3* using the *in vitro* translation experiment. The plasmid containing *ZmBTF3-TAA-ISR-Fluc* was firstly transcribed *in vitro*, followed by *in vitro* translation using wheat germ extract. *ZmBTF3* with TAA mutated to ACA was used as the positive control of the translation to measure the efficiency of SCR. The *Zm00001d009138-TAG-ISR-Fluc* was used as the SCR-negative control. Luciferase activity reflects the readthrough activity. All experiments were performed in triplicates. Data are mean ± s.e.m (n=3). Statistical difference was calculated using a two-sided Student’s *t*-test, \*\*\**P* < 0.001. ISR, inter-stop codon region. **c** SDS-PAGE and immunoblot analysis of the ZmBTF3x-Fluc (ZmBTF3-TAA-ISR-Fluc), ZmBTF3x^ACA^-Fluc (ZmBTF3-ACA-ISR-Fluc), and ZmBTF3-Fluc fusion proteins. Top: Coomassie Brilliant Blue (CBB)-stained gel of the expressed products derived from (**b**). Bottom: Immunoblot of the ZmBTF3x-Fluc, ZmBTF3x^ACA^-Fluc, and ZmBTF3-Fluc fusion proteins using an anti-Fluc antibody. The ZmBTF3x^ACA^-Fluc was used as the positive control. The Zm00001d009138-TAG-ISR-Fluc was used as the negative control. ISR, inter-stop codon region. **d** MS/MS analysis of the digested peptides of the ZmBTF3x-Fluc fusion protein excised from the CBB gel in (**c**). The sequences of MS-detected peptides, labeled as “before”, “through”, and “after”, are shown on the top. An MS/MS spectrum of the “through” peptide EPEEKKESTFCCV is shown on the bottom. UAA was recoded as threonine (T) shown in red. **e** Transmembrane domain prediction in the C-terminus of ZmBTF3x using the TMHMM server. The blue, red, and magenta lines represent residues 1-201 inside the membrane, 202-221 as a transmembrane helix, and 222-223 outside the membrane, respectively. The amino acid sequence from the annotated stop codon to the next stop codon of BTF3x is shown on the top. **f** AlphaFold structural models for ZmBTF3 and its readthrough protein ZmBTF3x. Green: ZmBTF3; dark blue: parent protein region of the readthrough protein ZmBTF3x; magenta: C-terminal extension region of ZmBTF3x. The predicted α-helix is shown at the C-terminus. **g** Subcellular localization of ZmBTF3 and ZmBTF3x. Constructs encoding C-terminal GFP fusions of ZmBTF3 or ZmBTF3x were transfected into maize protoplasts for localization assays under a confocal microscope. GFP-vector transfected protoplasts were used as the control. The experiments were performed three times with similar results. HY5-mCherry was the nucleus marker and PDCK7-mCherry was the plasma membrane marker. Scale bars, 5 μm. Source data are provided as a Source Data file.

The single transmembrane domain predicted in the C-terminus of ZmBTF3x was found to contain 20 amino acid residues crossing the membrane (Fig. 6e). The predicted 3D structures of ZmBTF3 and ZmBTF3x revealed an α-helix appended at the ZmBTF3x C-terminus that was not present in ZmBTF3 (Fig. 6f). Furthermore, the α-helix with a high pLDDT score and low PAE score corresponded exactly to the region of the predicted transmembrane domain (Fig. 6f and Supplementary Fig. 10a, b). To examine the biological significance of the C-terminal extension derived from SCR of *ZmBTF3*, we constructed *35S:ZmBTF3-GFP* and *35S:ZmBTF3x-GFP* vectors and individually transfected them in maize protoplasts for subcellular localization analysis using confocal microscopy. The results showed that ZmBTF3 localized to the nucleus and cytosol (Fig. 6g), consistent with the previous study reporting that *Arabidopsis* BTF3 acted as the transcription factor and the β-subunit of nascent polypeptide-associated complex (NAC)^53^. By contrast, the readthrough protein ZmBTF3x localized to the plasma membrane and cytosol (Fig. 6g), confirming the localization change between the isoforms ZmBTF3 and ZmBTF3x. In summary, these results indicate that SCR can lead to the structural and subcellular localization change of the readthrough protein isoform, which may confer new biological functions.

## Discussion

SCR, classified as the recoding of stop codons, is a regulatory mode of translation that could produce multiple protein isoforms from the same transcript^1, 54^. SCR has been reported in a variety of organisms with important biological implications. Recently, Sahoo *et al*. have identified mRNAs of 144 genes that exhibited SCR in *Arabidopsis* by analyzing Ribo-seq datasets^38^. However, a systematic study of SCR events in plants at the protein level is still lacking. In the current study, we employed a high-resolution MS-based proteogenomic strategy to analyze SCR events in different plants and to identify the amino acids translated at stop codon sites. We established 20 customized databases for each of the four plant species, which enabled detection of any potential SCR events with all possible amino acids translated at stop codon sites. Using this strategy, we detected 1,009 SCR events corresponding to a collection of 2,185 proteins in four plant species.

Our systematic analysis of SCR events revealed their similar characteristics among the four plant species. Firstly, SCR events occur in all chromosomes of the four species, and SCR events are prone to be dispersively distributed in the chromosome, except in maize where the higher distribution of SCR was closer to telomeres. Secondly, consistent with previous studies^6, 23^, UGA is the most preferential stop codon for readthrough, and it also turned out to be the most preferred next in-frame stop codon in SCR events of these species. In addition, SCR transcripts were mostly significantly shorter and had fewer exons and splice variants than non-SCR transcripts in two monocot plants, although these were not as significant as that in the dicots. However, in other eukaryotes like *D. melanogaster* and *Anopheles gambiae*, SCR transcripts were found to be much longer than the non-SCR transcripts, possibly because the occurrence of SCR requires more regulatory elements embedded in their longer transcripts^23, 24^. Our findings suggest that the regulation of SCR in plants might differ from that in animals.

The stop codons are able to be recoded as all 20 standard amino acids in the four plant species in our study, which has not been reported elsewhere. This result suggests a greater recoding potential of stop codons underestimated in the past and reveals a more flexible mode of decoding. Since tRNA is the adapter molecule in decoding a codon to an amino acid, we explored the reasons for the plasticity of stop codons by tRNA analysis. Suppressor tRNAs, of which the anti-codon could fully form canonical base-pairing with stop codons, have been reported to contribute to the translation of stop codons^24, 55, 56^. Here, we detected an array of suppressor tRNAs corresponding to different amino acids in maize, which may be the underlying reasons for the translation of six amino acids at UGA, six amino acids at UAG, and eight amino acids at UAA sites. In addition to suppressor tRNAs, near-cognate tRNAs of stop codons are also an important reason for the translation due to the noncanonical base pairing of one position between a stop codon and the anticodon of the tRNA^41, 44^. The amino acids carried by these near-cognate tRNAs include arginine, serine, tryptophan, cysteine, glycine, and leucine for UGA; lysine, glutamate, glutamine, tryptophan, serine, leucine, and tyrosine for UAG; and lysine, glutamate, glutamine, tyrosine, serine, and leucine for UAA^57, 58^. However, we did not find suppressor tRNAs for the rest small portion of the 20 amino acids recoded at stop codon sites in our study, nor the tRNAs of these amino acids were near-cognate tRNAs (e.g., the case of UAG recoded as alanine and UGA recoded as glutamate in Fig. 2). Indeed, these amino acids are not exclusively observed in this study, as previous studies have also reported some of their translation at stop codon sites^15, 30, 39, 46^. These cases suggest that our present understanding of stop codon reading by suppressor or near-cognate tRNAs is still incomplete. Considering the sophisticated modifications and regulation of the decoding adaptor tRNAs and multiple factors that influence SCR events^8, 59^, the mechanism underlying SCR associated with different tRNAs warrants further study.

In summary, we implemented a proteogenomic strategy and identified abundant SCR events across the genomes of four plant species. Stop codons in these SCR events could be recoded as all 20 standard amino acids. We demonstrated that SCR events are widespread in plant species and uncovered the unprecedented plasticity of stop codons. Our findings provide evidence that genetic decoding is far more flexible than has been recognized. We also identified a large volume of novel protein isoforms derived from SCR, which not only expands the plant proteome but also lays a foundation for further exploring the functionality of SCR products.

## Methods

### Sample preparation

Maize (*Zea mays,* B73 inbred line), rice (*Oryza sativa* L. ssp. *japonica* cv. Nipponbare), *Arabidopsis thaliana* (Columbia-0), and soybean (*Glycine max* L., ZH13) plants were grown in a temperature- and light-controlled greenhouse. The growth conditions for maize were a 15-h light/9-h dark photoperiod and a 28/25 ℃ day/night air temperature, and samples were collected at the three-leaf stage. For rice, the growth conditions were a 12-h light/12-h dark photoperiod and a 28/25 ℃ day/night air temperature, and samples were collected on day 56. *Arabidopsis* plants were grown in a greenhouse under a 16-h light/8-h dark photoperiod and a 22/21 ℃ day/night air temperature, and samples were collected on day 21. Soybean plants were grown in a greenhouse under a 16-h light/8-h dark photoperiod and a 22/21 ℃ day/night air temperature, until the trifoliate leaves were fully expanded. Three replicates of leaf samples for each of the plant species were harvested, frozen in liquid nitrogen, and stored at –80 ℃ until analysis.

### Protein extraction

Total proteins from each sample were extracted as previously reported^60^. Briefly, leaf tissues (1 g) for each plant species were pulverized in liquid nitrogen, and the powder was precipitated in 10% (w/v) trichloroacetic acid/acetone solution containing 65 mM dithiothreitol (DTT) at –20 °C for 1 h. Then, the solution was centrifuged at 10,000×g at 4 ℃ for 45 min and the supernatant was removed. The precipitate was then vacuum-dried and solubilized in 1/10 volumes of SDT buffer (4% [w/v] SDS, 100 mM DTT, and 150 mM Tris-HCl, pH 8.0). After a 3-min incubation in boiling water, the suspensions were ultrasonicated (80 w, ultrasonic 10 s/time, every 15 s, repeated 10 times) and incubated at 100 ℃ for 3 min. The extract was centrifuged at 12,000×g at 25 ℃ for 10 min, and the total protein contents were quantified using BCA Protein Assay Reagent (Promega Corporation). The supernatants were stored at –80 ℃ until subsequent analysis.

### Protein digestion and peptide fractionation

The total proteins (250 μg for each sample) were digested by FASP procedure using trypsin (Promega Corporation) as described previously^60^. Then, the digested peptide filtrate was fractionated by strong cation exchange (SCX) chromatography using an AKTA Purifier system (GE Healthcare). The dried peptide mixture was reconstituted and acidified with 2 ml buffer A (10 mM KH^2^PO^4^ in 25%[v/v] of ACN, pH = 2.7) and loaded onto a PolySULFOETHYL 4.6×100 mm column (5 µm, 200 Å, PolyLC Inc). The peptides were eluted at a flow rate of 1 ml/min buffer B (500 mM KCl, 10 mM KH^2^PO^4^ in 25% of ACN, pH 2.7) in a gradient of 0-10% (v/v) for 2 min, 10-20% (v/v) for 25 min, 20-45% (v/v) for 5 min, and 50-100% (v/v) for 5 min. The elution was monitored by absorbance at 214 nm, and fractions were collected every 1 min. The 30 collected fractions were combined into 15 pools and desalted on C18 Cartridges (Empore SPE Cartridges C18 standard density, bed I.D. 7 mm, volume 3 ml, Sigma-Aldrich). Each fraction was concentrated by vacuum centrifugation and reconstituted in 40 µl of 0.1% (v/v) trifluoroacetic acid. All samples were stored at –80 °C before LC-MS/MS analysis.

### Liquid chromatography-tandem mass spectrometry analysis

The samples were analyzed on a Q-Exactive HF-X mass spectrometer coupled to the Easy nLC 1200 (Thermo Fisher Scientific). Peptide (2 μg) was loaded onto a 25 cm, 75 μm inner diameter C18-reversed phase column packed with 3 μm resin (EASY-Column Capillary Columns, Thermo Fisher Scientific) in buffer A (2% [v/v] acetonitrile and 0.1% [v/v] formic acid) and separated with a linear gradient of buffer B (80% [v/v] acetonitrile and 0.1% formic acid) at a flow rate of 250 nl/min controlled by IntelliFlow technology over 90 min. MS data were acquired using a data-dependent top 10 method dynamically choosing the most abundant precursor ions from the survey scan (m/z 300-1,800) for higher-energy collision dissociation fragmentation (HCD). Determination of the target value was based on predictive Automatic Gain Control (pAGC). The dynamic exclusion duration was 25 s. Survey scans were acquired at a resolution of 70,000 (m/z 200), and the resolution for HCD spectra was set to 17,500 (m/z 200). Normalized collision energy was 30 eV, and the underfill ratio, which specifies the minimum percentage of the target value likely to be reached at maximum fill time, was defined as 0.1%. The instrument was run with peptide recognition mode enabled. MS experiments for each sample were conducted in triplicate for each sample.

### Customized database construction

The complete genome sequences of the four plants were downloaded from Ensembl Plants (ftp://ftp.ensemblgenomes.org/pub/plants/release-41/fasta/zea_mays/dna/) for maize, MSU Rice Genome Annotation Project (http://rice.uga.edu/pub/data/Eukaryotic_Projects/o_sativa/annotation_dbs/pseudomol ecules/version_7.0/all.dir/) for rice, the National Genomics Data Center (ftp://download.big.ac.cn/gwh/Plants/Glycine_max_Gmax_ZH13_v2.0_GWHAAEV 00000000.1/GWHAAEV00000000.1.genome.fasta.gz) for soybean, and TAIR 10 (ftp://ftp.ensemblgenomes.org/pub/plants/release-28/fasta/Arabidopsis_thaliana/dna/Arabidopsis_thaliana.TAIR10.28.dna.toplevel.fa.gz) for *Arabidopsis* in FASTA format. The 20 customized databases for each plant species were constructed as previously described with modifications^46^.

### Identification of SCR events by MaxQuant

The raw data were analyzed using MaxQuant software (v.1.6.1.10)^61^. The spectra were searched against the 20 customized databases for each of the four plant species. The initial search was set at a precursor mass window of 6 ppm. The search followed an enzymatic cleavage rule of trypsin and allowed a maximum of two missed cleavage sites and a mass tolerance of 20 ppm for fragment ions. Carbamidomethylation of cysteines was defined as fixed modification, while methionine oxidation was defined as variable modification for database searching. The cutoff of global FDR for peptide and protein identification was set to 0.01. Protein quantification was calculated by normalized spectral protein intensity.

All matched peptides were then relocated to the corresponding positions of protein sequences in the reference databases and mapped to their genomic locations. Every stop codon site was given an ID for genomic position recognition. The non-redundant peptides uniquely mapped to the regions spanning the annotated stop codon were labeled as “through”, the peptides mapped to the coding sequence (CDS) region were labeled as “before”, and the peptides mapped to the 3′ UTR were labeled as “after”. We further implemented more stringent criteria to determine the “through” peptides so as to identify SCR events. Only the unique “through” peptides that met the following conditions were retained: (1) presented in at least two of the three replicates in MS experiment; (2) at least two of the quality scores were higher than 20. Finally, proteins sharing one stop codon site and containing a unique “through” peptide were merged into one group and the group was defined as one SCR event.

### Ribosome profiling analysis

We obtained ribosome profiling (Ribo-seq) datasets of 56 maize samples from previous reports for SCR event analyzing^62–66^. The Ribo-seq data was analyzed according to the methods described by Dunn et al. (2013)^11^ and Sapkota et al. (2019)^6^ with modifications. Briefly, the ribosomal sequencing data were quality-controlled using fastax (v.36.06) or cutadapt (v.1.13)^62^ to remove low-quality bases, adaptor sequences, and random primers. Reads of 25-35 nucleotides (nts) were retained by seqkit (v.0.5.4) for subsequent analysis. The remaining reads were aligned to the maize genome by STAR (v.2.7.3a)^67^. Ribosome-protected footprint alignments were mapped to their estimated P-sites as follows: 12 nt were cut from each end of the alignment, resulting in a fragment of *n* × nt in length (*n* = *l*-12×2). Each genomic position covered by a nucleotide remaining in the truncated alignment was then incremented by 1/*n* (count number). Then, the reads were mapped to the following positions: 12 nts after the start codon, 15 nts before and 9 nts after the first stop codon, and 15 nts after the second stop codon. This resulted in four regions remaining in the transcripts: the 5′ UTR, CDS, readthrough region, and the distal 3′ UTR.

We then analyzed the length distribution of Ribo-seq reads in each dataset. Four Ribo-seq datasets in which the read length was mainly distributed in 30-nt (30-mer) and prepared with RNase I (little cutting bias) were finally retained for further SCR validation. For analysis of 3-nt periodicity (phasing) of footprints, R package ribosomeProfilingQC (v.1.10.0) was used and only 30-nt RPFs were counted, as shown in maize with good-phased footprint population^62^. Reads were then counted in all putative extended regions. Mean value of normalized RPKM (reads per kilobase per million mapped reads) was calculated in R (v.3.2.5) in the four datasets. The ribosome footprint density of putative extensions and the distal 3′ UTRs were calculated in RPKM. Only extensions with RPKM > 1, read coverage > 10%, and no overlap with any annotated 5′ UTRs, CDSs, and other non-coding RNAs were retained and regarded as readthrough. Full data of samples that exhibited readthrough in Ribo-seq analysis for each of the SCR events are shown in Supplementary Table 3.

### Distribution of readthrough events at the genomic level

The genome-wide density of SCR and non-SCR events in maize chromosomes was plotted using a sliding window of 20 Mb with 2-Mb steps according to the annotated maize genome in Ensembl (https://plants.ensembl.org/index.html). We used “hotspot region” to reflect the distribution density of SCR and non-SCR events. Hotspot regions were defined as ≥ 5 SCR events for SCR hotspots and ≥ 335 for non-SCR events in a window size of 20 Mb. The chromosomal distributions of rice SCR and non-SCR events were drawn using a window size of 4 Mb with a step of 400 kb based on the annotated rice genome from MSU (http://rice.uga.edu/). The same distribution pattern was plotted with a window size of 10 Mb with a 1-Mb step based on the ZH13 reference genome (https://ngdc.cncb.ac.cn/gwh/Assembly/652/show) for soybean. For *Arabidopsis*, the SCR and non-SCR genomic localizations were obtained by a 1.2-Mb sliding window with a step size of 100 kb according to the annotated *Arabidopsis* genome in TAIR (https://www.Arabidopsis.org/). Hotspot regions were defined as ≥3 for SCR events and ≥178 for non-SCR events in a window size of 4 Mb for rice, ≥2 for SCR events and ≥253 for non-SCR events in a window size of 10 Mb for soybean, and ≥4 for SCR events and ≥157 for non-SCR events in a window size of 1.2 Mb for *Arabidopsis*.

### Motif and minimum free energy analysis in readthrough regions of SCR transcripts in maize

Transcription factor-binding motifs in readthrough regions of maize SCR transcripts were identified using the MEME-ChIP webserver (v.5.4.1)^68^, and the annotations of the obtained motifs were searched against the available databases for comparison using Tomtom (v.5.0.5) with default parameters^69^.

The minimum free energy of readthrough regions for maize SCR transcripts and the comparable length between the first and downstream second stop codons for non-readthrough transcripts were calculated using ViennaRNA Package (v.2.0) as described^70^.

### Identification and alignment of suppressor tRNAs in maize

The complete maize genome sequences were downloaded from the Ensembl database (ftp://ftp.ensemblgenomes.org/pub/plants/release-41/fasta/zea_mays/dna/). A genome-wide search for suppressor tRNA genes was conducted using tRNAscan-SE (v.2.0)^51^ with default settings and ARAGORN^52^ with both default settings and 0.95 default settings as a lower scoring threshold. This analysis generated a collection of suppressor/suppressor-like tRNAs recognizing the three stop codons from each search. Comparison of the canonical tRNAs from different species with these predicted suppressor tRNAs was performed using BLASTn (v.2.5.0+) against the NCBI GenBank nucleotide (NCBI NT) database as previously described^55, 56^. The potential identity of the suppressor tRNAs corresponding to different amino acids could be predicted from the high similarity of the sequence alignment between the suppressor tRNAs and the canonical tRNAs of the standard amino acids.

### Motif and structure prediction of SCR proteins in maize

C-terminal extensions at least 20 amino acids in length in maize SCR events were scanned for transmembrane helices using TMHMM (v.2.0)^71^ with default settings and for nuclear localization signals using cNLS mapper^72^. C-terminal extensions at least 12 amino acids in length were searched for peroxisomal targeting signals using PTS1 Predictor^73^ and prenylation signals using PrePS^74^. The 3D structures of ZmBTF3 and ZmBTF3x were predicted using Alphafold^75^.

### Plasmid construction

Luciferase constructs for the luminescence-based SCR assay were generated in the reconstructive pFN19K T7 SP6 flexi vector (Promega Corporation). The coding sequences of the tested genes, *ZmH3C2*, *ZmCEQORH*, *ZmBTF3* and *Zm00001d009138*, along with their canonical stop codons and inter-stop codon regions (ISRs, the extension regions) were cloned upstream of the coding sequence of the firefly luciferase gene (*Fluc*) without ATG between *EcoRI* and *KpnI* sites. A linker sequence (5′ GGCGGCTCCGGCGGCTCCCTCGTGCTCGGG 3′) was added upstream of the Fluc sequence to avoid potential distortions between the tested protein and Fluc in the fusion protein^15, 31, 38^. Primer sequences are listed in Supplementary Table 15.

For subcellular localization analysis, the ZmBTF3 and ZmBTF3x coding sequences were introduced into the pAN580 vector containing the 35S promoter and the coding sequence for GFP. GFP was fused to the C-terminus of ZmBTF3 and ZmBTF3x. Primer sequences are listed in Supplementary Table 15.

### *In vitro* translation for SCR Assay

For SCR assay, we used *in vitro* translation system based on Wheat Germ Extract (Promega Corporation), which has been used for SCR validation as reported previously^31, 38^. The constructed plasmid DNA including the test gene and *Fluc* (no ATG) was linearized using BamH I or XbaI enzyme, and 2 μg of the linearized DNA was transcribed *in vitro* using T7 RNA polymerase (Promega Corporation). The obtained RNA was then purified using GeneJET kit (Thermo Fisher Scientific). The concentration and purity of the RNA were measured using Nanodrop 2000 (Thermo Fisher Scientific). A total of 2 μg purified RNA was translated *in vitro* using the Wheat Germ Extract at 25 °C for 2 h according to the manufacturer′s instructions. Luciferase activity was then measured at room temperature (below 25 °C) using the Luciferase Assay System (Promega Corporation) in a Glomax luminometer (Promega Corporation) with a 10-s measurement read for luciferase activity.

### Immunoblotting and mass spectrometric analysis

The proteins extracted from the *in vitro* protein expression system were separated by 10% (w/v) SDS-polyacrylamide gel electrophoresis (SDS-PAGE). One PAGE gel was stained with Coomassie Brilliant Blue (CBB), while another was transferred to a polyvinylidene difluoride membrane (0.22 µm, Millipore). Proteins were detected by immunoblotting with anti-Fluc antibodies (1:10,000, ab185924, Abcam). The regions of interest in the CBB staining gel were excised manually according to the immunoblotting band, and then destained with 30% (v/v) ACN in 100 mM NH^4^HCO^3^ and dried in a vacuum centrifuge. The extracted proteins were reduced with dithiothreitol (DTT) (10 mM DTT in 100 mM NH^4^HCO^3^) for 30 min at 56 °C, then alkylated with iodoacetamide (200 mM IAA in 100 mM NH^4^HCO^3^) in the dark at room temperature for 30 min. Gel pieces were briefly rinsed with 100 mM NH^4^HCO^3^ and ACN, respectively. The gel pieces were then divided equally into three parts and separately digested overnight in 12.5 ng/μl trypsin, overnight in chymotrypsin in 25 mM NH^4^HCO^3^, and for 30 min in 12.5 ng/μl pepsin in PH3 HCl. The peptides were extracted using 60% (v/v) ACN in 0.1% (v/v) trifluoroacetic acid (TFA) three times, and the extracts were pooled and dried completely by a vacuum centrifuge.

LC-MS/MS was performed on a Q-Exactive HF-X mass spectrometer coupled to Easy nLC 1200 (Thermo Fisher Scientific). The peptide mixture was loaded onto a C18-reversed phase column (15 cm, 75 μm inner diameter) packed in-house with RP-C18 (5 μm resin, Thermo Fisher Scientific) in buffer A (0.1% [v/v] formic acid in HPLC-grade water) and separated with a linear gradient of buffer B (0.1% formic acid in 84% [v/v] ACN) at a flow rate of 250 nl/min controlled by IntelliFlow technology over 60 min. MS data were acquired using a data-dependent top-10 method dynamically choosing the most abundant precursor ions from the survey scan (m/z 300-1,800) for HCD fragmentation. The target value was determined based on pAGC. The dynamic exclusion duration was 20 s. Survey scans were acquired at a resolution of 70,000 (m/z 200) and the resolution for HCD spectra was set to 17,500 (m/z 200). Normalized collision energy was 27 eV and the underfill ratio, which specifies the minimum percentage of the target value likely to be reached at maximum fill time, was defined as 0.1%.

The MS data were then separately searched against the target protein sequences (readthrough protein sequences of *H3C2*, *ZmCEQORH*, and *ZmBTF3*) using MaxQuant software (v.1.6.1.10)^61^. An initial search was set with a precursor mass window of 6 ppm. The search followed an enzymatic cleavage rule of “unspecified” and a mass tolerance of 20 ppm for fragment ions. Carbamidomethylation of cysteines was defined as fixed modification, while methionine oxidation was defined as variable modification for database searching. The cutoff of the global FDR for peptide and protein identification was set to 0.01.

### Subcellular localization of ZmBTF3-GFP and ZmBTF3x-GFP fusions

The GFP fusion plasmids were transformed into maize mesophyll protoplasts by polyethylene glycol-mediated transformation. Protoplast preparation and transfection were conducted as previously described^76^. A plasma membrane-localized marker *35S:ZmPDCK7-mCherry* and a nucleus-localized marker *35S:AtHY5-mCherry* were used for colocalization analyses^77, 78^. Fluorescence signals were observed on a confocal laser scanning microscope (LSM800, Zeiss).

## Data availability

All data generated or analyzed in this study are included in this article or the Supplementary Information and Tables. The raw datasets of proteomics have been deposited at the ProteomeXchange under the accession ID PXD035958 and PXD038762. Source data are provided.

## Code availability

Custom codes are available for research purposes from the corresponding authors upon request.

## Acknowledgments

This study was supported by the National Natural Science Foundation of China (no. 32172073, U22A20474, 31872872), the National Key Research and Development Program of China (2022YFD1201802), and the Seed-Industrialized Development Program in Shandong Province (no. 2021LZGC003). We are grateful to Steven P. Briggs and David Backhouse for their constructive comments and suggestions. We also thank Dr. Anguo Sun and Dr. Yanwen Xiang for the technical assistance and Yingying Sha and Mengxia Niu for their assistance with the microscopic observations.

## Author Contributions

L.W., S.-B.W., and Z.T. designed the project. Y. Zhang, Y.S., L.T., H.Y., A.X., J.Z., U.A., A.S., X.C. and Y.C. were involved in bioinformatics analysis. Y. Zhang, H.L., S.W. and J.S. conducted the experiments. C.G., Y. Zhao, Y. L. and X.W. collected the materials for mass spectrometry detection and subsequent experiments. Y. Zhang, H.L. and Y.S. wrote the manuscript. L.W., S.-B.W., and Z.T. contributed substantially to revisions. All authors read and approved the manuscript.

## Supplementary Information, Tables and Dataset have been submitted as separate files

**Supplementary Information contains Supplementary** Figures 1-10 **and References for Supplementary Table 5**

**Supplementary Table 1. SCR events identified in maize by proteogenomics.**

**Supplementary Table 2. Peptides identified by mass spectrometry for SCR and parent proteins in maize.**

**Supplementary Table 3. SCR events detected in maize by both proteogenomics and ribosome profiling.**

**Supplementary Table 4. Non-SCR events identified in maize by proteogenomics.**

**Supplementary Table 5. Standard amino acids translated by stop codons in the literature and their translation evidence in maize. I, UGA; II, UAG; III, UAA. UGA-, UAG-, and UAA suppressor tRNAs were predicted by tRNAscan-SE or ARAGORN.**

**Supplementary Table 6. Predicted suppressor tRNAs in maize aligned with tRNAs in the NCBI NT database.**

**Supplementary Table 7. SCR events identified in rice by proteogenomics.**

**Supplementary Table 8. Non-SCR events identified in rice by proteogenomics.**

**Supplementary Table 9. SCR events identified in soybean by proteogenomics.**

**Supplementary Table 10. Non-SCR events identified in soybean by proteogenomics.**

**Supplementary Table 11. SCR events identified in *Arabidopsis* by proteogenomics.**

**Supplementary Table 12. Non-SCR events identified in *Arabidopsis* by proteogenomics.**

**Supplementary Table 13. Functional signals predicted in C-terminal extensions of maize SCR events.**

**Supplementary Table 14. Primers used in this study.**

**Supplementary Dataset 1: Motifs predicted by Tomtom in the C-terminal extensions of SCR transcripts in maize.**

