## Supplementary Fig. for "Widespread readthrough events in plants reveal unprecedented plasticity of stop codons"

**for**

Zhang *et al*

**Supplementary Figures 1-10**

**Supplementary References**

### Supplementary Figures

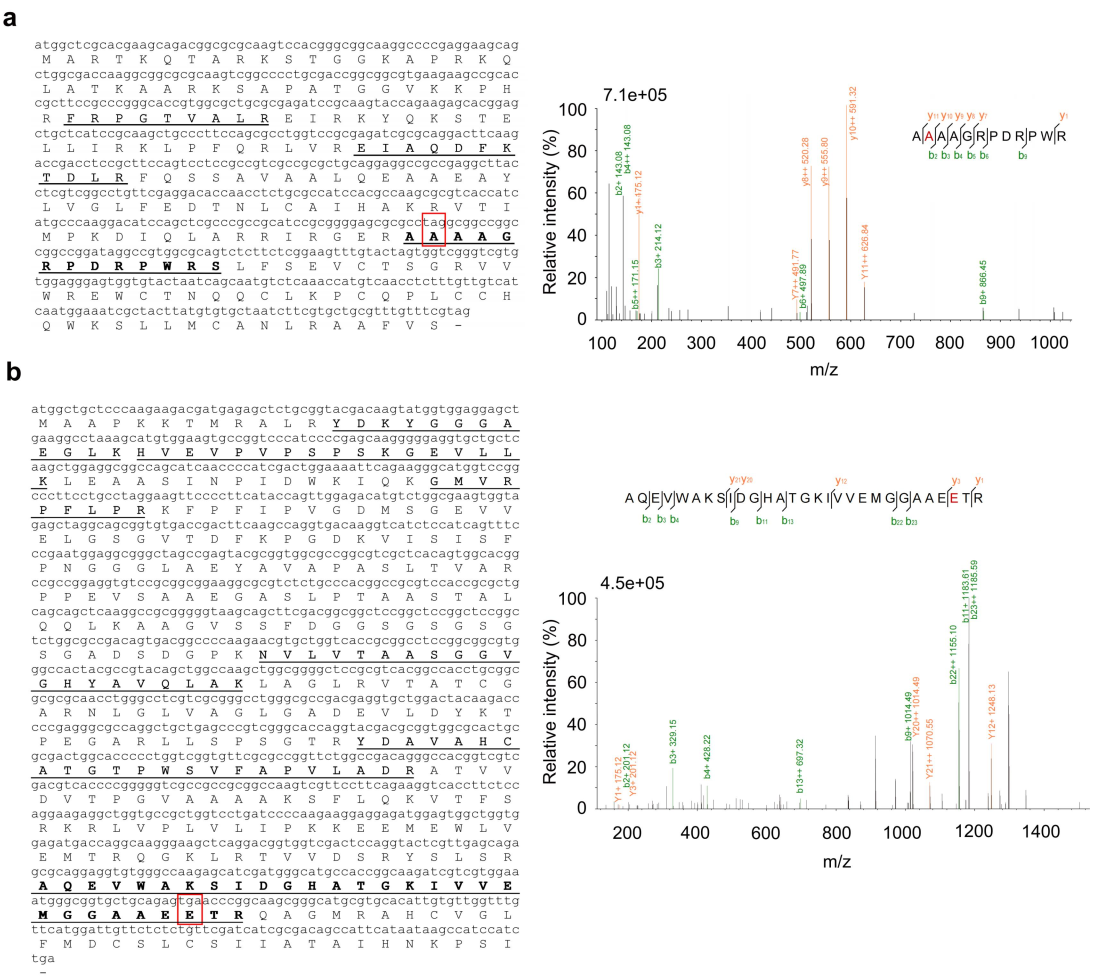


**Supplementary Fig. 1. Peptides identified by mass spectrometry and MS/MS spectra of readthrough peptides for two examples of SCR in maize. a, b,** Left: peptides identified by mass spectrometry mapped onto the extended protein sequences translated to the next in-frame stop codons of *ZmH3C2* (**a**) and *ZmCEQORH* (**b**). Amino acids translated at different stop codon sites are boxed in red, which are alanine (A) at UAG (**a**) and glutamate (E) at UGA (**b**). Right: MS/MS spectra of readthrough peptides "AAAAGRPDRPWR" for ZmH3C2x (**a**) and "AQEVWAKSIDGHATGKIVVEMGGAAEETR" for ZmCEQORHx (**b**).

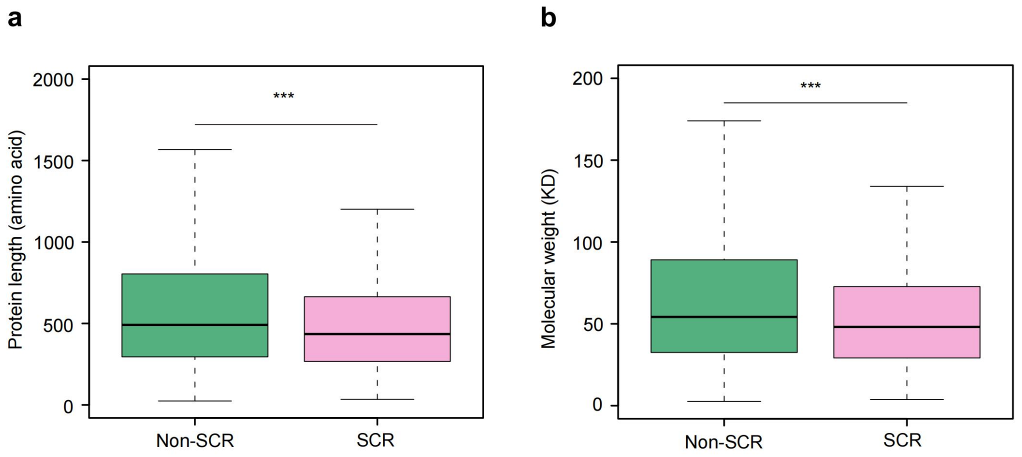


**Supplementary Fig. 2. Characteristics of proteins in SCR events in maize. a,** Amino acid length of proteins in SCR and non-SCR events. **b,** Molecular weight of proteins in SCR and non-SCR events. For all boxplots, box edges represent the 0.25 and 0.75 quantiles. The bold lines represent the median values and the whiskers represent 1.5×IQR. Statistical difference was determined using a two-sided Student's *t*-test, ***$P < 0.001$.

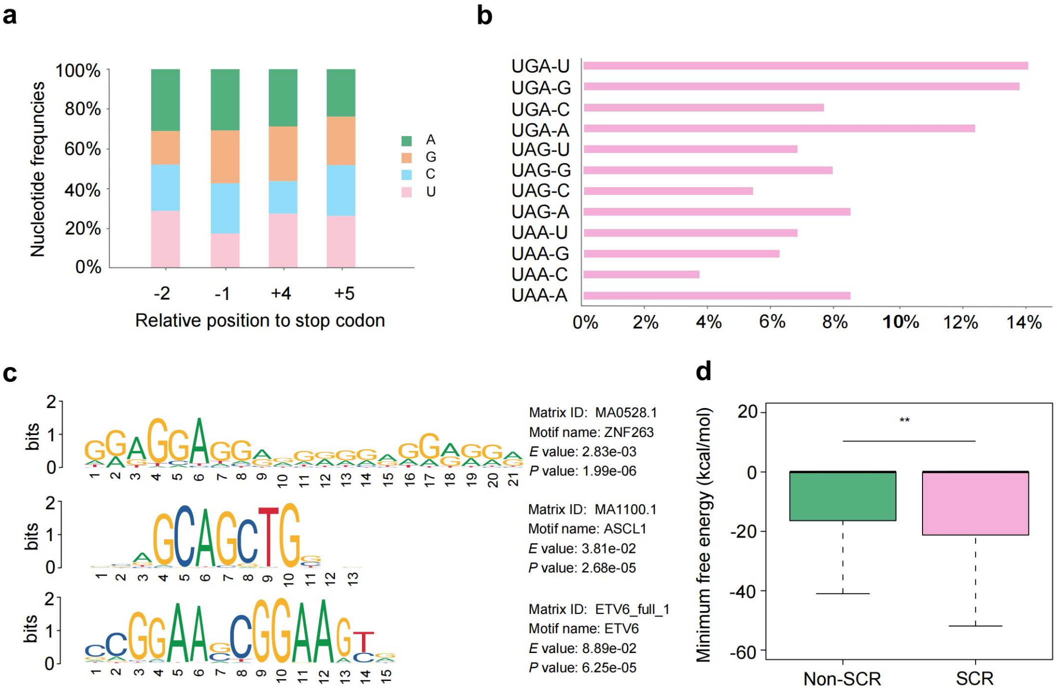


**Supplementary Fig. 3. Local mRNA context of SCR transcripts in maize. a,** Nucleotide usage in positions -2, -1, +4, and +5 immediately preceding and following the readthrough stop codons. **b,** Tetranucleotide pattern distribution (stop codon plus the subsequent nucleotide) in maize SCR events. **c,** Three representative motifs located in the readthrough regions of SCR transcripts. **d,** Comparison of the minimum free energy of the regions from the annotated stop codon to the next in-frame stop codon between SCR and non-SCR transcripts. For the boxplot, box edges represent the 0.25 and 0.75 quantiles. The bold lines represent the median values and the whiskers represent 1.5×IQR. A two-sided Student's *t*-test was used to calculate the statistical difference, $^{**}P < 0.01$.

**a**

```
tRNA-(uca)_[152291517152291600]

                                  (((((((..((((.........))))(((((.......))))).)................
tRNA-(uca)                        GGGGUGGUGGCGCAGUUGGCUAGCGCGUAGGUCUCACAUCUG-UGAGUGAUCCUGAGGUC  59
Oryza sativa_AJ506175.1_Met       GGGGUGGUGGCGCAGUUGGCUAGCGCGUAGGUCUCAUAGCUAUUGAGUGAUCCUGAGGUC  60
Oryza sativa_AJ506176.1_Met       GGGGUGGTGGCGCAGUUGGCUAGCGCGUAGGUCUCAUAGCUAUUGAGUTAUCCUGAGGUC  60
Glycine max_M12150.1_Met          GGGGUGGUGGCGCAGUUGGCUAGCGCGUAGGUCUCAUA-----------AUCCUGAGGUC  49
Arabidopsis thaliana_D50934.1_Met GGGGUGGUGGCGCAGUUGGCUAGCGCGUAGGUCUCAUA-----------AUCCUGAGGUC  49
Arabidopsis thaliana_D50933.1_Met GGGGUGGUGGCGCAGUUGGCUAGCGCGUAGGUCUCAUA-----------AUCCUGAGGUC  49
Arabidopsis thaliana_D50932.1_Met GGGGUGGUGGCGCAGUUGGCUAGCGCGUAGGUCUCAUA-----------AUCCUGAGGUC  49
Triticum aestivum_K02003.1_Met    GGGGUGGUGGCGCAGUUGGCUAGCGCGUAGGUCUCAUA-----------AUCCUGAGGUC  49
                                  *********************************** *          ***********

                                  (((((.......))))))))))))).
tRNA-(uca)                        GAGAGUUCGAGCCUCUCUCACCCCA--------     84
Oryza sativa_AJ506175.1_Met       GAGAGUUCGAGCCUCTCUCACCCC---------     84
Oryza sativa_AJ506176.1_Met       GAGAGUUCGAGCCUCUCUCACCCC---------     84
Glycine max_M12150.1_Met          GAGAGUUCGAGCCUCUCUCACCCCA--------     74
Arabidopsis thaliana_D50934.1_Met GAGAGUUCGAGCCUCUCUCACCCCA--------     74
Arabidopsis thaliana_D50933.1_Met GAGAGUUCGAGCCUCUCUCACCCCA--------     74
Arabidopsis thaliana_D50932.1_Met GAGAGUUCGAGCCUCUCUCACCCCA--------     74
Triticum aestivum_K02003.1_Met    GAGAGUUCGAGCCUCUCUCACCCCACCA-----     77
                                  ************************
```

**b**

```
tRNA-(uca)_[107765514107765586]

                                 (((((((...(((.........))).((((((.......))))))).....((((......
tRNA-(uca)                       AGCCCAUCUAGCCUAGUUGGUAGACCACAAGGCUUCAAACCUUAUGGUCGUGGGUUCAAG   60
Trypanosoma brucei_X57047.1_lys  -GCCCUUCUAGCUCAGUCGGUAGAGCGCACGGCUCUUAACCGUGUGGUCGUGGGUUCGAG   59
Trypanosoma brucei_X57044.1_lys  UGCCCUUCUAGCUCAGUCGGUAGAGCGCACGGCUCUUAACCGUGUGGUCGUGGGUUCGAG   60
Trypanosoma brucei_S57044.1_lys  -GCCCUUCUAGCUCAGUCGGUAGAGCGCACGGCUCUUAACCGUGUGGUCGUGGGUUCGAG   59
Trypanosoma brucei_X57045.1_Lys  --GCCUUCCUAGCUCAGUGGUAGAGCGCACGGCUUUUAACCGUGUGGUCGUGGGUUCGAU   58
Trypanosoma brucei_Z11880.1_Lys  --GCCUUCCUAGCUCAGUGGUAGAGCGCACGGCUUUUAACCGUGUGGUCGUGGGUUCGAU   58
                                    ** **          ****** * ** ****   **** * ************* *

                                 ))))))))))))..
tRNA-(uca)                       CCCCAUAGUUUAC- 73
Trypanosoma brucei_X57047.1_lys  CCCCACGGGGGGUG 73
Trypanosoma brucei_X57044.1_lys  CCCCACGGGGGGUG 74
Trypanosoma brucei_S57044.1_lys  CCCCACGGGGGGUG 73
Trypanosoma brucei_X57045.1_Lys  CCCCACGGAAGGCG 72
Trypanosoma brucei_Z11880.1_Lys  CCCCACGGAAGGCG 72
                                 *****
```

**Supplementary Fig. 4. Nucleotide sequence alignments of different UGA suppressor tRNAs in maize with tRNAs in the NCBI NT database. a,** Alignments of UGA suppressor tRNAs with $tRNAs^{Met}$. **b,** Alignments of UGA suppressor tRNAs with $tRNAs^{Lys}$.

**a**

```
tRNA-(cua)_[2645006526450137]|7.trna16_7:26450065-26450137_(+)_Sup_(CUA)

                                    ((((((((..((((.........))))((((((.......))))))...(((((.......
tRNA-(cua)                          UCCGUUGUAGUCUAGCUGGUUAGGAUCCUCGGCUCUAAUCCGAGAGACCCAGGUUCGAGU  60
Lupinus luteus_M23387.1_Glu         UCCGUCGUAGUCUAGGDGGDUAGGAUACUCGGCUUUCACCCGAGAGACCCGGGUUCAAGU  60
Hordeum vulgare_M22136.1_Glu        UCCGUCGUAGUCUAGGDGGDUAGGAUACUCGGCUUUCACCCGAGAGACCCGGGUUCAAGU  60
Arabidopsis thaliana_AB005786.1_Glu UCCGUUGUAGUCUAGCUGGUCAGGAUACUCGGCUCUCACCCGAGAGACCCGGGUUCGAGU  60
Zea mays_KJ526738.1_Glu             UCCGUUGUAGUCUAGCUGGUCAGGAUAUUCGGCUCUCACCCGAAAGACCCGGGUUCGUGU  60
                                    ***** *********  **  *****  ****** * * **** ****** *****  **

                                    ))))))))))))).
tRNA-(cua)                          CCCGGCAACGGAA---    73
Lupinus luteus_M23387.1_Glu         CCCGGCGACGGAACCA    76
Hordeum vulgare_M22136.1_Glu        CCCGGCGACGGAACCA    76
Arabidopsis thaliana_AB005786.1_Glu CCCGGCAACGGAG---    73
Zea mays_KJ526738.1_Glu             CCCGGCAACGGAACCA    76
                                    ****** *****
```

**b**

```
tRNA-(cua)_c[257655602257655673]|1.trna172_1:257655602-257655673_(-)_Sup_(CUA)

                                  ((((((((..((((.......))))((((((.......))))))....(((((.......)
tRNA-(cua)                        GAAUCUGUGGCGUAAUGGUAGCGCGUCUGACUCUAGAUCAGAAGGUUGAGUGUAUGAUUC  60
Homo sapiens_HG984064.1_Trp       GGCCUCGUGGCGCAACGGUAGCGCGUCUGACUCCAGAUCAGAAGGUUGCGUGUUCAAAUC  60
Homo sapiens_HG984070.1_Trp       GACCUCGUGGCGCAACGGCAGCGCGUCUGACUCCAGAUCAGAAGGUUGCGUGUUCAAAUC  60
Homo sapiens_HG984981.1_Trp       GACCUUGUGGCGCAACGGUAGCGCGUCUGACUCCAGAUCAGAAGGUUGCGUGUUCAAAUC  60
Homo sapiens_HG984066.1_Trp       GACCUCGUGGCGCAACGGUAGCGCGUCUGACUCCAGAUCAGAAGGUUGCGUGUUCAAAUC  60
Avian sarcoma virus_M17490.1_Trp  GACCUCGUGGCGCAACGGDAGCGCGUCUGACUCCAGAUCAGAAGGCUGCGUGUUCGAAUC  60
Homo sapiens_HG984069.1_Trp       GACCUCGUGGCGCAACGGUAGCGCGUCUGACUCCAGAUCAGAAGGCUGCGUGUUCGAAUC  60
Avian oncornavirus_M10671.1_Trp   GACCUCGUGGCGCAACGGUAGCGCGUCUGACUCCAGAUCAGAAGGCUGCGUGUUCGAAUC  60
Gallus gallus_X07245.1_Trp        -ACCUCGUGGCGCAACGGUAGCGCGUCUGACUCCAGAUCAGAAGGCUGCGUGUUCGAAUC  59
Homo sapiens_HG984564.1_Trp       GACCUCGUGGCACAAUGGUAGCACGUCUGACUCCAGAUCAGAAGGUUGCGUGUUCAAAUC  60
Bos taurus_K00264.1_Trp           GACCUCGUGGCGCAAUGGUAGCGCGUCUGACUCCAGAUCAGAAGGUUGCGUGUUCGAAUC  60
Homo sapiens_HG984557.1_Trp       GACCUCGUGGCGCAAUGGUAGCGCGUCUGACUCCAGAUCAGAAGGUUGCGUGUUCAAAUC  60
Homo sapiens_HG984065.1_Trp       GACCUCGUGGCGCAAUGGUAGCGCGUCUGACUCCAGAUCAGAAGGUUGCGUGUUCAAGUC  60
Nicotiana rustica_X64326.1_Trp    GGAUCCGUGGCGCAAUGGUAGCGCGUCUGACUCCAGAUCAGAAUGUUGCGUGUUCCAUUC  60
Arabidopsis thaliana_X57592.1_Trp GGAUCCGUGGCGCAAUGGUAGCGCGUCUGACUCCAGAUCAGAAGGUUGCGUGUUCGAUUC  60
Arabidopsis thaliana_L34745.1_Trp GGAUCCGUGGCGCAAUGGUAGCGCGUCUGACUCCAGAUCAGAAGGUUGCGUGUUCGAUUC  60
Arabidopsis thaliana_L35907.1_Trp GGAUCCGUGGCGCAAUGGUAGCGCGUCUGACUCCAGAUCAGAAGGUUGCGUGUUCGAUUC  60
Arabidopsis thaliana_L35908.1_Trp GGAUCCGUGGCGCAAUGGUAGCGCGUCUGACUCCAGAUCAGAAGGUUGCGUGUUCGAUUC  60
Arabidopsis thaliana_L35909.1_Trp GGAUCCGUGGCGCAAUGGUAGCGCGUCUGACUCCAGAUCAGAAGGUUGCGUGUUCGAUUC  60
Arabidopsis thaliana_X57594.1_Trp GGAUCCGUGGCGCAAUGGUAGCGCGUCUGACUCCAGAUCAGAAGGUUGCGUGUUCGAUUC  60
Arabidopsis thaliana_X57593.1_Trp GGAUUCGUGGCGCAAUGGUAGCGCGUCUGACUCCAGAUCAGAAGGUUGCGUGUUCGAUUC  60
                                        *****  ** ** *** ********** ********* * ** ****   * **

                                  ))))))))))))).
tRNA-(cua)                        ACGUCGGGUUCA------------------------------------    72
Homo sapiens_HG984064.1_Trp       ACGUCGGGGUCA------------------------------------    72
Homo sapiens_HG984070.1_Trp       ACGUCGGGGUCA------------------------------------    72
Homo sapiens_HG984981.1_Trp       ACGUCGGGGUCA------------------------------------    72
Homo sapiens_HG984066.1_Trp       ACGUCGGGGUCA------------------------------------    72
Avian sarcoma virus_M17490.1_Trp  ACGUCGGGGUCACCA---------------------------------    75
Homo sapiens_HG984069.1_Trp       ACGUCGGGGUCA------------------------------------    72
Avian oncornavirus_M10671.1_Trp   ACGUCGGGGUCACCA---------------------------------    75
Gallus gallus_X07245.1_Trp        ACGUCGGGGUCA------------------------------------    71
Homo sapiens_HG984564.1_Trp       ACGUCGGGGUCA------------------------------------    72
Bos taurus_K00264.1_Trp           ACGUCGGGGUCACCA---------------------------------    75
Homo sapiens_HG984557.1_Trp       ACGUCGGGGUCA------------------------------------    72
Homo sapiens_HG984065.1_Trp       ACGUCGGGGUCA------------------------------------    72
Nicotiana rustica_X64326.1_Trp    ACGUCGGGUUCACCA---------------------------------    75
Arabidopsis thaliana_X57592.1_Trp ACGUCGGGUUCAAAUAUCCCGA--------------------------    82
Arabidopsis thaliana_L34745.1_Trp ACGUCGGGUUCAAACCCCCGGUUUAAACAUGAUCCUUAACUUUUUAUUU    109
Arabidopsis thaliana_L35907.1_Trp ACGUCGGGUUCAAUCCCCCGAACAAUUUCCGGUUUUUUUUUUUUUUC--    107
Arabidopsis thaliana_L35908.1_Trp ACGUCGGGUUCAAAUCCCCCGAACAAUAAAUUCCGGAAUAUUUUUUU--    107
Arabidopsis thaliana_L35909.1_Trp ACGUCGGGUUCAAAUCCCCCGAACUAUAUAUUCCGGAAUAU--------    101
Arabidopsis thaliana_X57594.1_Trp ACGUCGGGUUCA------------------------------------    72
Arabidopsis thaliana_X57593.1_Trp ACGUCGGGUUCA------------------------------------    72
                                  ******** ***
```

**Supplementary Fig. 5. Nucleotide sequence alignments of UAG suppressor tRNAs in maize with tRNAs in the NCBI NT database. a,** Alignments of UAG suppressor tRNAs with $tRNAs^{Glu}$. **b,** Alignments of UAG suppressor tRNAs with $tRNAs^{Trp}$.

**a**

```
tRNA-(uua)_[67391246739196]|9.trna10_9:6739124-6739196_(+)_Sup_(UUA)

                                     ((((((..((((........)))).(((((.......))))).....(((((......
tRNA-(uua)                           -GCACGUCUAGCUCAGUCGGUAGAGCGCAAGACUUUAAACCUUGUGGUCAUAGGUUUGAG      59
Heterololigo bleekeri_D50538.1_Lys   -GCCCGGCUAGCUCAGUCGGUAGAGCACGAGACUUUUAAUCUCGGGGUCGUGGGUUCAAG      59
Dictyostelium discoideum_X59581.1_Lys  -GCCCAGAUAGCUCAGUCGGUAGAGCGCAAGGCUUUUAACCUUGUGGUCGGGGGUUCGAG      59
Dictyostelium discoideum_X59580.1_Lys  -GCCCAGAUAGCUCAGUCGGUAGAGCGCAAGGCUUUUAACCUUGUGGUCGGGGGUUCGAG      59
Dictyostelium discoideum_X59579.1_Lys  -GCCCAGAUAGCUCAGUCGGUAGAGCGCAAGGCUUUUAACCUUGUGGUCGGGGGUUCGAG      59
Dictyostelium discoideum_X59578.1_Lys  -GCCCAGAUAGCUCAGUCGGUAGAGCGCAAGGCUUUUAACCUUGUGGUCGGGGGUUCGAG      59
Dictyostelium discoideum_X59577.1_Lys  -GCCCAGAUAGCUCAGUCGGUAGAGCGCAAGGCUUUUAACCUUGUGGUCGGGGGUUCGAG      59
Dictyostelium discoideum_X59576.1_Lys  -GCCCAGAUAGCUCAGUCGGUAGAGCGCAAGGCUUUUAACCUUGUGGUCGGGGGUUCGAG      59
Dictyostelium discoideum_X59575.1_Lys  -GCCCGGCUAGCUCAGUCGGUAGAGCGCCAGACUCUUAAUCUGGUGGUCGGGGGUUCGAG      59
Dictyostelium discoideum_X59574.1_Lys  -GCCCGGCUAGCUCAGUCGGUAGAGCGCCAGACUCUUAAUCUGGUGGUCGGGGGUUCGAG      59
Trypanosoma brucei_X57047.1_Lys      -GCCCUUCUAGCUCAGUCGGUAGAGCGCACGGCUCUUAACCGUGUGGUCGUGGGUUCGAG      59
Trypanosoma brucei_X57044.1_Lys      UGCCCUUCUAGCUCAGUCGGUAGAGCGCACGGCUCUUAACCGUGUGGUCGUGGGUUCGAG      60
Trypanosoma brucei_S57044.1_Lys      -GCCCUUCUAGCUCAGUCGGUAGAGCGCACGGCUCUUAACCGUGUGGUCGUGGGUUCGAG      59
                                      ** *   ***************** *  * ** * ** *  * ****   ****  **

                                     .))))))))))))).
tRNA-(uua)                           CCCCACGGUGGGCG-  73
Heterololigo bleekeri_D50538.1_Lys   CCCCACGUUGGGCU-  73
Dictyostelium discoideum_X59581.1_Lys  CCCCCCUUUGGGCG-  73
Dictyostelium discoideum_X59580.1_Lys  CCCCCCUUUGGGCG-  73
Dictyostelium discoideum_X59579.1_Lys  CCCCCCUUUGGGCG-  73
Dictyostelium discoideum_X59578.1_Lys  CCCCCCUUUGGGCG-  73
Dictyostelium discoideum_X59577.1_Lys  CCCCCCUUUGGGCG-  73
Dictyostelium discoideum_X59576.1_Lys  CCCCCCUUUGGGCG-  73
Dictyostelium discoideum_X59575.1_Lys  CCCCCCGUUGGGCG-  73
Dictyostelium discoideum_X59574.1_Lys  CCCCCCGUUGGGCG-  73
Trypanosoma brucei_X57047.1_Lys      CCCCACGGGGGGUG-  73
Trypanosoma brucei_X57044.1_Lys      CCCCACGGGGGGUG-  74
Trypanosoma brucei_S57044.1_Lys      CCCCACGGGGGGUG-  73
                                     **** *   ***
```

**b**

```
tRNA-(uua)_[2890195428902027]

                                          (((((((...(((.(........)))).(((((.......)))))......((((.....
tRNA-(uua)                                AAGCCAUCUAGCU-CAGUUGGUGGAGCACAAGGUUUUAAACCAUGAAGUUGGGGGUUCGA  59
Desulfotalea psychrophila_LK006174.1_His  -GUGGGCGUGGCUCAGUUGGUUAGAGCACCUGGUUGUGACCCAGGAAGUCGGGGGUUCGA  59
                                                  * ***    * * * **** *  **** * * *** ***** **********
                                          .))))))))))))...
tRNA-(uua)                                GCCCUAUAACUUGCC---    74
Desulfotalea psychrophila_LK006174.1_His  GUCCCCUCGCUCACCCCA    77
                                          * **  *   ** **
```

**Supplementary Fig. 6. Nucleotide sequence alignments of UAA suppressor tRNAs in maize with tRNAs in the NCBI NT database. a,** Alignments of UAA suppressor tRNAs with tRNAs$^{Lys}$. **b,** Alignments of UAA suppressor tRNAs with tRNAs$^{His}$.

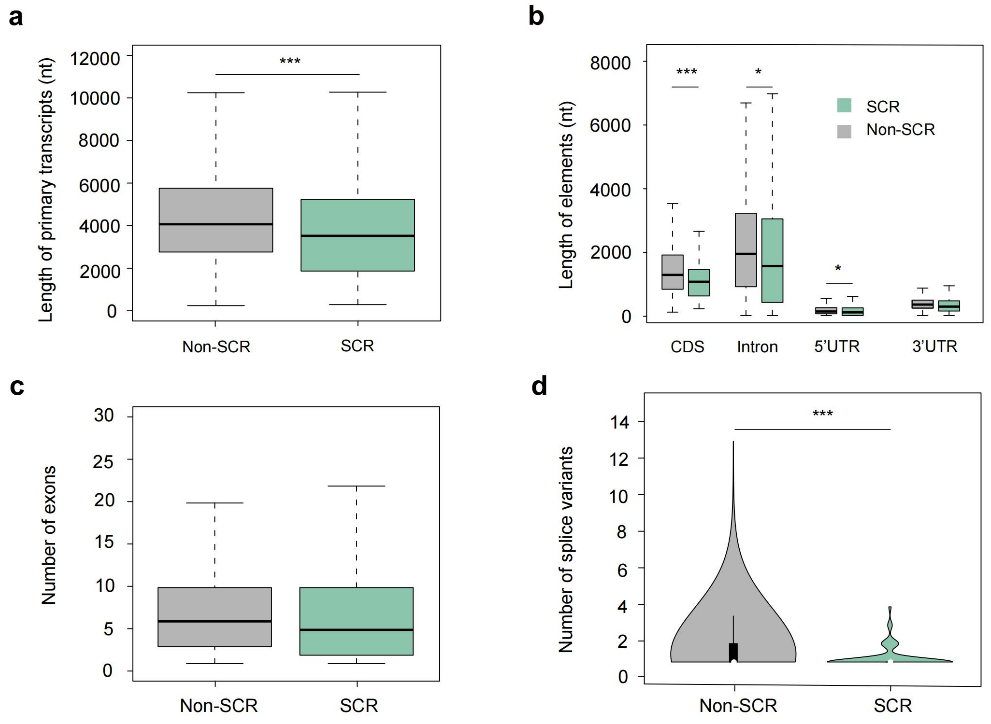


**Supplementary Fig. 7. Features of SCR Events in rice. a,** Length of SCR and non-SCR primary transcripts. The length of the primary transcript is the sum of CDSs, introns, and UTRs. **b,** Length of transcript elements of SCR and non-SCR transcripts. CDS length is the length of the ORF. **c,** Number of exons of SCR and non-SCR primary transcripts. **d,** Violin plot showing the number of splice variants of SCR and non-SCR primary transcripts. For all boxplots, box edges represent the 0.25 and 0.75 quantiles. The bold lines represent the median. The whiskers represent 1.5×IQR. For the violin plot, white dots in the internal boxplot represent the median values. Statistical difference was calculated using a two-sided Student's *t*-test, **P* < 0.05, ****P* < 0.001.

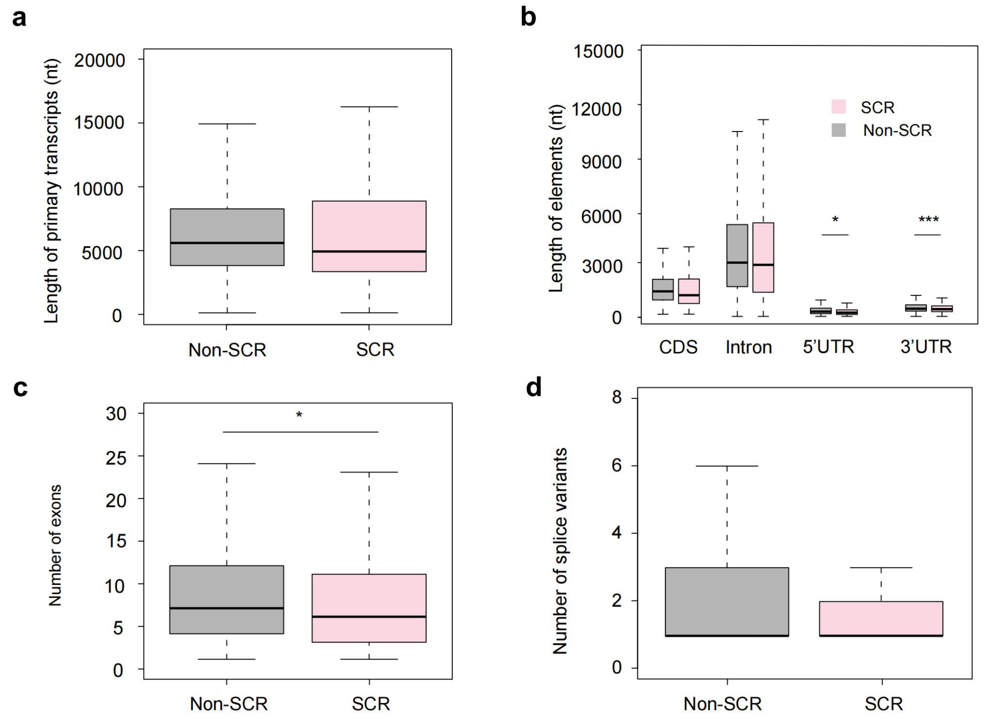


**Supplementary Fig. 8. Features of SCR transcripts in soybean. a,** Length of SCR and non-SCR primary transcripts. The length of the primary transcript is the sum of CDSs, introns, and UTRs. **b,** Length of transcript elements in SCR and non-SCR primary transcript. CDS length is the length of the ORF. **c,** Number of exons in SCR and non-SCR primary transcripts. **d,** Number of splice variants in SCR and non-SCR primary transcripts. For all boxplots, box edges represent the 0.25 and 0.75 quantiles. The bold lines represent the median values. The whiskers represent 1.5×IQR. Statistical difference was calculated using a two-sided Student's *t*-test, *$P < 0.05$, ***$P < 0.001$.

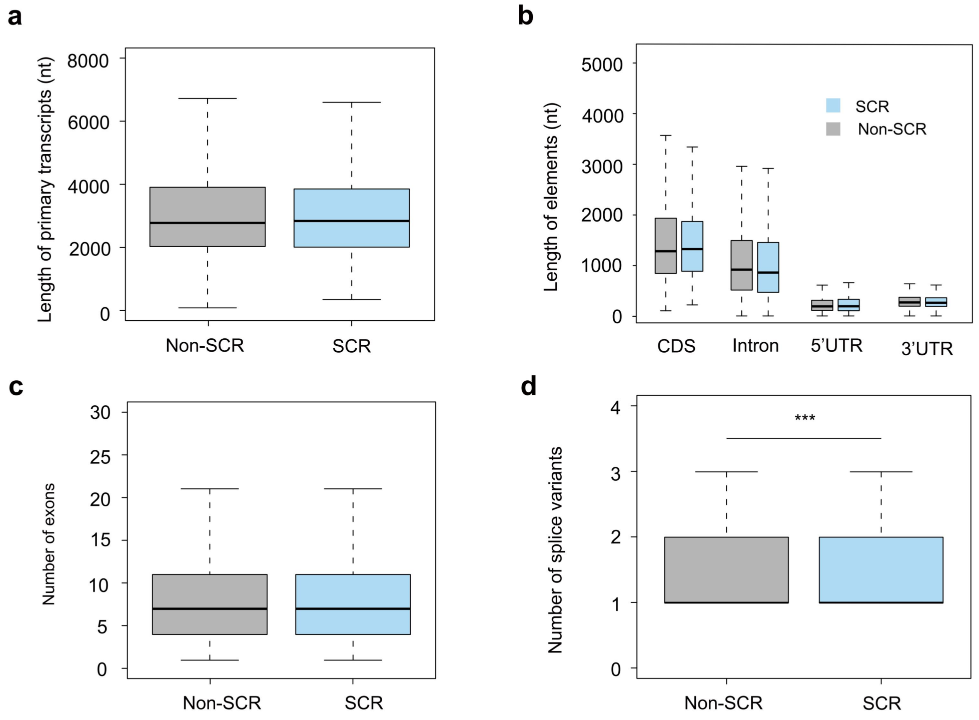


**Supplementary Fig. 9. Features of SCR transcripts in *Arabidopsis*. a,** Length of SCR and non-SCR primary transcripts. The length of the primary transcript is the sum of CDSs, introns, and UTRs. **b,** Length of transcript elements in SCR and non-SCR primary transcript. CDS length is the length of the ORF. **c,** Number of exons in SCR and non-SCR primary transcripts. **d,** Number of splice variants in SCR and non-SCR primary transcripts. For all boxplots, box edges represent the 0.25 and 0.75 quantiles. The bold lines represent the median values. The whiskers represent 1.5×IQR. Statistical difference was calculated using two-sided Student's *t*-test, ****P* < 0.001.

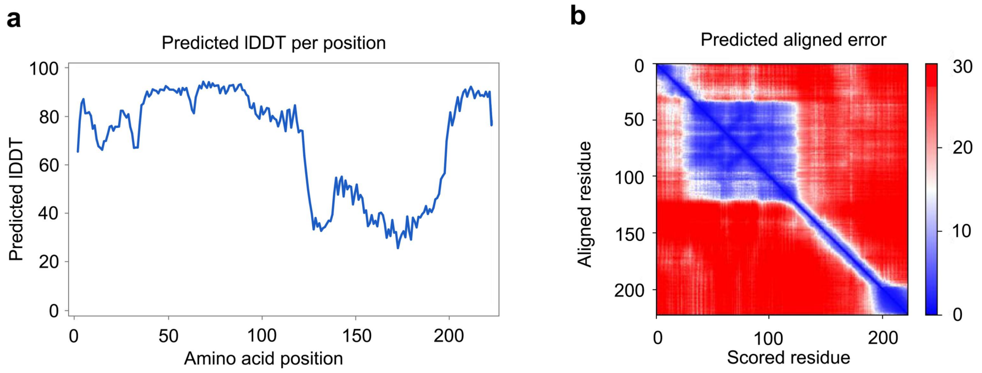


**Supplementary Fig. 10. AlphaFold metrics of 3D structure prediction for the readthrough protein ZmBTF3x. a,** The predicted lDDT (pLDDT) score is plotted per residue. lDDT is to assess the local accuracy of the prediction, awarding a high score for regions that are well-predicted. pLDDT is a confidence metric based on lDDT which reflects local confidence in the structure to assess confidence within a single domain. AlphaFold produces pLDDT score between 0 and 100: very high (pLDDT > 90); confident (90 > pLDDT > 70); low (70 > pLDDT > 50), and very low (pLDDT < 50). The predicted model has high confidence levels of pLDDT at the C-terminal α-helix region. **b,** Interactive 2D plot of Predicted Aligned Errors (PAEs). PAE indicates a distance error for every pair of residues, and the colour at (x, y) indicates the expected distance error in residue x's position, when the prediction and true structure are aligned on residue y. The blue color at (x, y) indicates that AlphaFold predicted low error with well-defined relative positions of different domains.
